# Ketamine restores escape behavior by re-engaging dopamine systems to drive cortical spinogenesis

**DOI:** 10.1101/2020.03.11.987818

**Authors:** M Wu, S Minkowicz, V Dumrongprechachan, P Hamilton, L Xiao, Y Kozorovitskiy

**Affiliations:** Department of Neurobiology, Northwestern University, Evanston, IL 60208

## Abstract

Escaping aversive stimuli is essential for complex organisms, but prolonged exposure to stress leads to maladaptive learning. Stress alters plasticity, neuromodulatory signaling, and neuronal activity in distributed networks, yet the field lacks a unifying framework for its varied consequences. Here we describe neuromodulatory and plasticity changes following aversive learning by using a learned helplessness paradigm, where ketamine restores escape behavior. Dopaminergic neuron activity in the ventral tegmental area systematically varies across learning, correlating with future sensitivity to ketamine treatment. Ketamine’s effects are blocked by chemogenetic inhibition of dopamine signaling and mimicked by optogenetic activation. We use 2-photon glutamate uncaging/imaging to interrogate structural plasticity in medial prefrontal cortex, revealing that dendritic spinogenesis on pyramidal neurons is both regulated by aversive experience and recovered by ketamine in a dopamine-dependent manner. Together, these data describe recurrent circuits that causally link neuromodulatory dynamics, aversive learning, and plasticity enhancements driven by a therapeutically promising antidepressant.

---

Ketamine and its S-enantiomer esketamine demonstrate rapid onset and lasting antidepressant effects in clinical trials^1,2^. They act primarily as antagonists of the glutamatergic N-methyl-D-aspartate (NMDA) receptors^3–5^, although some studies implicate mechanisms beyond direct NMDAR antagonism^6,7^. At the circuit level, emerging evidence supports a model of cortical disinhibition, where ketamine preferentially decreases synaptic activation of local inhibitory neurons, facilitating cortical activity^4,8,9^. Previous studies have shown that ketamine ameliorates depressive-like behaviors in animal models following acute or prolonged stress^10–14^. Accumulating evidence implicates the enhancement of synaptic plasticity in ketamine’s behavioral effects^5,12,13,15–18^. However, it is currently unknown which neural circuit dynamics mediate the effects of this rapidly acting antidepressant on behavior and plasticity.

Changes in reward-based and aversive learning represent features of depressive disorders that can be modeled in rodents^19–21^. One established model of aversive learning is learned helplessness^22–24^. Following prolonged inescapable stress exposure, animals learn that outcomes are independent of their behavioral actions; this learning eventually diminishes attempts to escape from avoidable stressful stimuli^25^. Specific activity patterns of dopaminergic (DA) neurons in the ventral tegmental area (VTA), and downstream DA release, encode reward and aversion^26–30^. This activity is differentially modulated by acute and chronic stress^31–34^, regulating depressive-like behaviors^35–37^. A recently published meta-analysis suggests that acute sub-anesthetic doses of ketamine may increase DA levels in the cortex, dorsal striatum, and nucleus accumbens^38^. Yet, a causal relationship between the DA system and ketamine’s effects on neural plasticity and behavior remains to be elucidated.

## VTA DA neuron activity during aversive learning

To define the function of midbrain DA neurons during aversive learning, we used a modified model of learned helplessness (LH)^5,32^. A shuttle box with two compartments connected by a door allows animals to escape from one side to the other when an electric foot shock is delivered to either compartment. Initially, mice escape from electric foot shocks. However, following repeated exposure to *inescapable* foot shocks, mice reduce escapes from avoidable 10 sec long foot shocks (Figure 1a, b). Our data and prior publications^34,40,41^ show that a single low dose of racemic ketamine 4 hrs prior to the test (10 mg/kg, b.w., i.p.) is sufficient to rescue escape behavior in this aversive learning paradigm (Figure 1b). In contrast, a separate group of mice that received saline instead of ketamine did not change the proportion of successful escapes after learning (Extended Data Figure 1a). Reduced escape behavior after LH induction, as well as its reversal by ketamine, are not strictly context-dependent, since the proportion of failures to escape was similar regardless of whether behavioral evaluation was carried out in the induction context or in a novel environment (Extended Data Figure 1b). Despite similar failure rates, differences in freezing behavior after LH induction were observed between environments prior to ketamine treatment (Extended Data Figure 1c). Similar failure rates in the induction and novel environments suggest a dissociation between escape propensity and freezing behavior.

**Figure 1.**
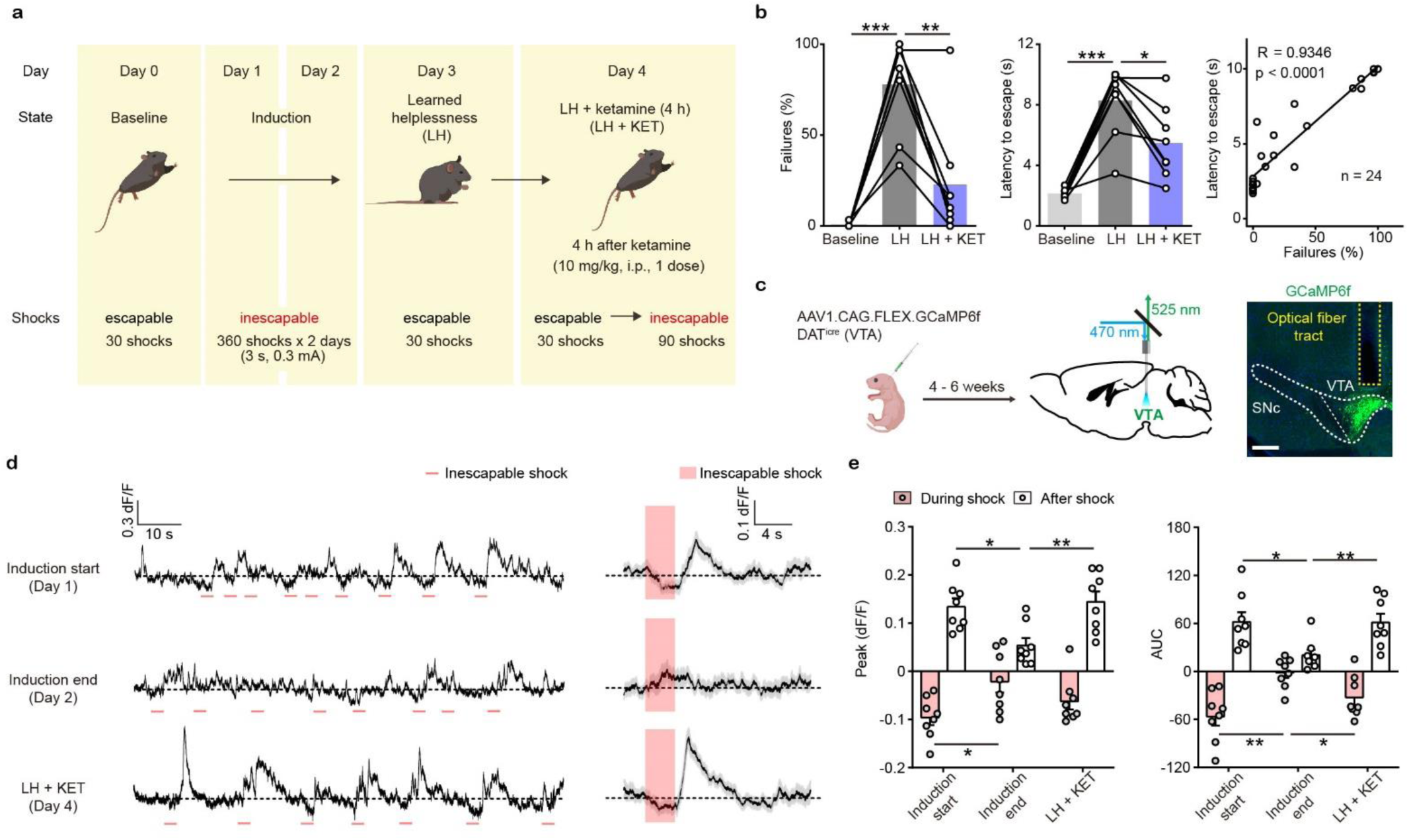
Ketamine rescues escape behavior and dampened DA neuronal activity after aversive learning. (a). Schematic illustrating the timeline of behavioral and pharmacological manipulations in the LH paradigm. (b). Left, summary data showing the percentage of failures to escape an escapable aversive shock across phases of learning (Baseline, LH, and LH + KET). Middle, same as left, but for latency to escape. Right, for all conditions, the correlation between percentage of failure to escape and escape latency (failure to escape trials scored as 10 sec latency). n = 24 trials from 8 mice. % Failures: repeated measures one-way ANOVA, F (1.89, 13.23) = 27.9, p = 0.0001, Sidak’s multiple comparison test, Baseline vs LH, p = 0.0001, LH vs LH+KET, p = 0.0037. Latency to escape: repeated measure one-way ANOVA, F (1.96, 13.72) = 29.63, p < 0.0001, Sidak’s multiple comparison test, Baseline vs LH, p = 0.0003, LH vs LH+KET, p = 0.0141. Pearson correlation: R = 0.9346, p < 0.0001. (c). Left, schematic for viral transduction in the VTA and subsequent fiber implant. Right, fiber placement verification. Green, GCaMP6f; blue, Hoechst nucleic stain. Scale bar: 500 μm. (d). Left, baseline adjusted raw traces of VTA DA neuron Ca^2+^ responses to inescapable foot shocks (3 sec, pink) in one animal, at the start of induction, at the end of induction, and 4 hrs following a single dose of ketamine (LH + KET, 10 mg/kg i.p.). Right, average traces in the same subject aligned to shock start time (20 trials/condition, mean ± SEM). (e). Left, quantification of peak Ca^2+^ transient amplitude during and after foot-shock stimuli across conditions. Right, same but for area under the curve (AUC). Both positive and negative values are quantified. n = 8 animals, repeated measures one-way ANOVA, Holm-Sidak’s multiple comparison test, Peak: During shock, F (1.823, 12.76) = 5.387, p = 0.0222, Induction start vs Induction end, p = 0.0458, Induction end vs LH+KET, p = 0.0788. After shock, F (1.693, 11.85) = 6.805, p = 0.0132, Induction start vs Induction end, p = 0.0230, Induction end vs LH+KET, p = 0.0068. AUC: During shock, F (1.705, 11.94) = 8.942, p = 0.0054, Induction start vs Induction end, p = 0.0058, Induction end vs LH+KET, p = 0.0258. After shock, F (1.437, 10.06) = 5.499, p = 0.0318, Induction start vs Induction end, p = 0.0257, Induction end vs LH+KET, p = 0.0069. *p < 0.05, ** p < 0.01, *** p< 0.001. Error bars reflect SEM.

To understand the relationship between VTA DA neuron activity and ketamine’s rescue of escape behavior, we used fiber photometry to monitor the activity of the genetically encoded calcium indicator GCaMP6f in VTA DA neurons. DAT^icre^ neonates were virally transduced with Cre-dependent AAV1.CAG.FLEX.GCaMP6f. Four to six weeks after transduction, optical fibers were implanted in the VTA (Figure 1c), guided by real-time photometry (Extended Data Figure 1d). The activity of VTA DA neurons was evaluated during aversive learning in young adult mice (P40-60) of both sexes. Surgical anesthesia during fiber implant procedure was associated with ∼1 Hz oscillations in VTA calcium transients (Extended Data Figure 1e). We first recorded Ca^2+^ transients of VTA DA neurons in response to brief, inescapable foot shocks (3 sec, 0.3 mA) during the learning period and after ketamine treatment. At the start of learning (Induction start), the activity of VTA DA neurons first decreased during the aversive foot shocks and then rose after the termination of the shock (Figure 1d, e). This biphasic response is consistent with recently published observations of VTA DA activity during other forms of aversive conditioning^42^. At the end of the second-day induction (Induction end), the responses of VTA DA neurons to inescapable foot shocks were blunted. A single low dose of racemic ketamine (10 mg/kg, b.w., i.p., LH+KET) largely restored the characteristic Ca^2+^ transient features, in parallel to the behavioral rescue (Figure 1d, e; Extended Data Figure 1g, h). Visualizing sequential Ca^2+^ traces from individual trials illustrates (1) the prevalence of after-shock peaks at the start of learning, (2) their decreased latency relative to shock onset during the second day of training, and (3) the recovery of Ca^2+^ transient latency to peak following ketamine administration (Extended Data Figure 1i). To quantify this temporal structure, we computed the latency to peak on sequential averaged trials (n = 10 trials/avg) and plotted the data as a scatter plot and cumulative distribution (Extended Data Figure 1j). No significant transients were observed in GFP-expressing controls in this behavioral assay (Extended Data Figure 1f).

Since the activity of DA neurons has been linked to movement^43,44^, one potential explanation for how aversive learning could modulate Ca^2+^ activity of VTA DA neurons involves altered locomotor behavior. While ketamine acutely changes locomotion, this effect normally takes places within tens of minutes following administration^9^, and resolves by the time clinically relevant changes in affective behavior are observed^2,45^. We found no differences in open field locomotion 4 hrs after ketamine injection across phases of learning (Extended Data Figure 1k). Additionally, changes in VTA DA neuron activity were not associated with motion transitions, including onset and offset of locomotion (Extended Data Figure 1l), a result that is not surprising, since movement transitions are typically associated with the activity of DA neurons in SNc^43,44^. These experiments demonstrate that VTA DA neuron activity is restructured by aversive learning, rather than by changes in locomotion.

To better understand the relationship between VTA DA neuron activity and specific behavioral responses, we designed a weaker learning paradigm (wLH), which allowed us to compare Ca^2+^ transients across distinct behavioral outcomes, escapes versus failures. wLH includes a large number of brief (3 sec long) escapable foot shocks as the test stimuli, with only a single day of LH induction with inescapable shocks (Figure 2a). As anticipated, a weaker form of learning, characterized by a lower average failure rate, was observed in this paradigm compared to stronger LH (sLH, as in Figure 1) (Two-way ANOVA for sLH and wLH across behavioral states, Sidak’s multiple comparison, sLH 78% vs wLH 48%, p = 0.0022). Again, escape behavior rapidly recovered with ketamine treatment (Figure 2b). Control animals underwent test sessions but no induction; they showed no changes in escape behavior over the time of the experiment (one-way ANOVA, p = 0.9506).

**Figure 2.**
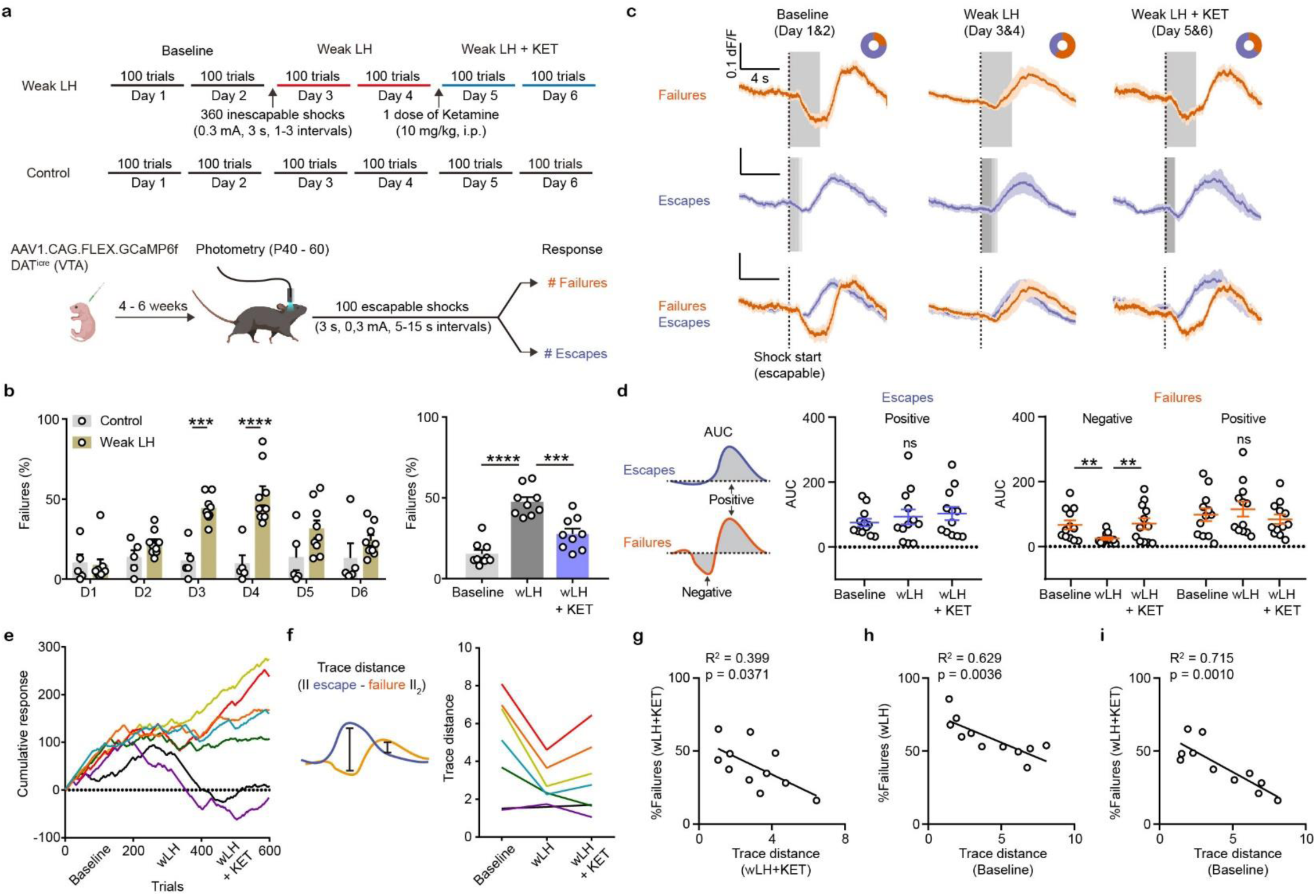
Weak learning analysis links DA activity, behavioral outcomes, and response to ketamine. (a). Top, schematic of experimental timeline for weak LH (wLH). Bottom, schematic of viral transduction, test trial description, and timing of photometry recording. (b). Left, summary data showing the percentage of failures to escape an escapable aversive shock across 6 days for two groups (white bar, controls; beige, wLH). Right, within group summary of behavioral data for wLH mice across learning phases. Control, n = 5, wLH, n = 9 animals. Left, responses across days, two-way ANOVA, Sidak’s multiple comparison test, Control vs wLH, D3, p = 0.0002, D4, p < 0.0001, D5, p = 0.103, D1, D2, and D6, p > 0.6. Right, repeated measures one-way ANOVA, F (1.908, 15.26) = 66.15, p < 0.0001. Sidak’s multiple comparison test, Baseline vs wLH, p < 0.0001, wLH vs wLH + KET, p = 0.0001. (c). Fiber photometry recordings of VTA DA Ca^2+^ transients, separated by behavioral response and aligned to shock start time (purple, escape; orange, failure; average dF/F across animals). Grey rectangles mark shock length in time, which is constant for failures to escape but variable for successful escapes and is shaded proportionally. Donut plots depict proportion of behavioral responses. n = 8 animals, baseline: 1405 trials, 25% failure 75% escape; wLH: 1459 trials, 60% failure 40% escape; wLH + KET: 1403 trials, 36% failure, 64% escape. (d). Left, schematic illustration for measured variables. Middle, summary data for AUC of the positive Ca^2+^ transient peaks in escapes across learning phases. Right, same but AUC for both positive and negative peaks in failures, as shown in the schematic. n = 12 animals, AUC (Escapes), positive peak, repeated measures one-way ANOVA, F (1.897, 20.87) = 1.881, p = 0.1787. AUC (Failures), negative peak, repeated measures one-way ANOVA, F (1.852, 20.38) = 9.260, p = 0.0017, Holm-Sidak’s multiple comparison test, Baseline vs wLH, p = 0.0069, wLH vs wLH + KET, p = 0.0069. Positive peak, repeated measures one-way ANOVA, F (1.540, 16.94) = 2.541, p = 0.1181. (e). Learning curves for individual animals. n = 7 animals, 600 trials/animal. (f). Left, schematic illustration for the measured variable, Euclidean norm of the difference between mean escape and failure-associated Ca^2+^ transients for each subject. Right, trace distances for each subject across learning phases with subject specific colors as in e. (g). Correlation between trace distance and the percentage of failures in wLH + KET. n = 11 animals. (h). Correlation between trace distance in the baseline condition and the percentage of failures in wLH. n = 11 animals. (i). Correlation between trace distance in the baseline condition and the percentage of failures in wLH + KET. n = 11 animals. ** p < 0.01, *** p < 0.001, **** p < 0.0001. Error bars reflect SEM.

A three-state transition model depicts the probability of transitioning between behavioral responses, allowing us to compare patterns of response sequences across learning (Extended Data Figure 2a). In addition to escapes and failures, responses were labeled ‘spontaneous’ when the animal spontaneously ran to the other side of the chamber during the random length pretrial time. Baseline and post ketamine sequences of behavioral responses were more similar to each other than either was to wLH. In addition to changing the average probabilities of pairs of responses, wLH learning also altered longer sequences of outcomes within animals, reflected in decreased probability and length of successive escapes, as well as increased successive failures. Ketamine treatment partially recovered the probability and length of successive escapes, while a more prominent effect was observed on decreasing successive failures back to baseline levels (Extended Data Figure 2b). Altogether, these analyses suggest that ketamine specifically restructures behavioral sequences that are altered by aversive learning.

The weaker learning along with a large number of escapable foot shocks as the test stimuli enabled broad sampling of Ca^2+^ transients during both escape and failure trials across learning. Similar reductions in VTA DA responses to inescapable foot shocks were observed during wLH induction as for strong LH (Extended Data Figure 2c). For escapable shocks, separately plotting VTA DA neuron activity for trials where the animal escaped or failed to escape revealed that each behavioral response is associated with distinct Ca^2+^ transient shapes (Figure 2c). Failure trial transients were biphasic, where fluorescence decreased during the shock and increased afterwards. In contrast, monophasic transients accompanied escape trials, with an increase in fluorescence following successful transitions to the other side of the box.

After learning, an increase in the similarity of the activity patterns between escape and failure trials was observed. For successful escape responses, VTA DA Ca^2+^ transients did not change between wLH and wLH + ketamine. The increased similarity of Ca^2+^ transients between outcomes after learning was explained by the reduction in negative transients associated with failures. These failure-linked negative transients recovered following ketamine treatment (Figure 2d). The timing of peaks in failure outcomes varied across learning conditions (Extended Data Figure 2d). We also noted a modest increase in overall fluorescence after termination of behavioral testing that emerged during learning (Extended Data Figure 2e).

In humans and other animals, individual variability in stress susceptibility and in responsiveness to neuroactive compounds is broadly acknowledged^46^. To visualize behavioral trajectories, learning curves were constructed by starting at 0 and incrementing by 1 for every escape trial, decrementing by 1 for every failure, and keeping constant for spontaneous transitions (Figure 2e). These learning curves highlight individual variability in learning and ketamine responsiveness. To depict the relationship between an animal’s ketamine responsiveness and VTA DA activity, we plotted the Euclidean norm of the difference between their Ca^2+^ transients during escape and failure trials (||*escape* − *failure*||_2_) across learning (Figure 2f, Extended Data Figure 2f). We selected this trace distance metric because it is agnostic to amplitude and kinetics of the response, specifically assaying the degree to which the activity of VTA DA neurons differentiates specific trial outcomes. Figure 2g shows a correlation between behavioral response (% failures) and trace distance following ketamine. Stronger ketamine behavioral effects were linked to more distinct Ca^2+^ transients when comparing escape and failure trials (R^2^ = 0.399). Animals characterized by a larger trace distance in the baseline were more resilient to inescapable stress during aversive learning, illustrated by fewer failures to escape after wLH (Figure 2h, R^2^ = 0.629). Intriguingly, individuals with larger differences between DA calcium signals on escape and failure trials in the baseline showed stronger responses to future ketamine treatment (Figure 2i, R^2^ = 0.715).

## DA neuron activity is necessary for behavioral effects of ketamine

The strong association between DA transients, behavioral states, and trial outcomes raises the possibility that particular signatures of DA signals in response to aversive stimuli are required for escape actions. To test whether VTA DA activity is required for escape actions in general, or alternatively, whether it becomes important for restoring escapes after learning, we conditionally expressed an inhibitory DREADD (designer receptor exclusively activated by designer drug) hM4Di in VTA DA neurons. This engineered muscarinic receptor is activated by clozapine-N-oxide (CNO) to drive Gα_i_-coupled pathways^47–50^. We used a CBA promoter driven Cre-conditional hM4Di AAV, with expression in VTA DA neurons confirmed by immunofluorescence (Figure 3a). Efficacy was evaluated using cell-attached recordings of hM4Di-mCherry^+^ VTA DA neurons, with a flow in and a washout of 1 µM CNO (Figure 3b).

**Figure 3.**
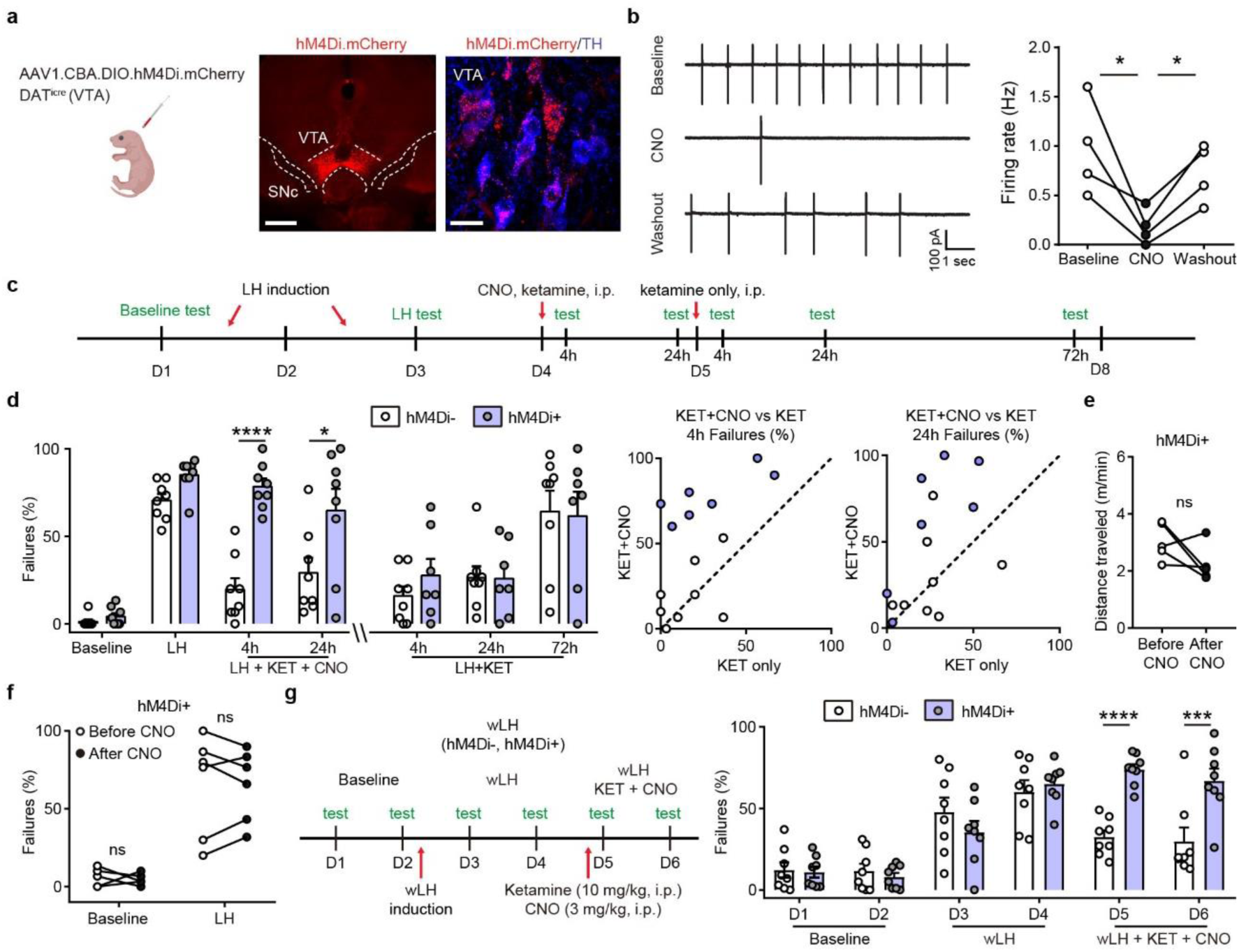
Ketamine behavioral effects in aversive learning require VTA DA activity. (a). Schematic for viral transduction in the VTA and hM4Di expression in VTA DA neurons (red, hM4Di.mCherry; blue, tyrosine hydroxylase (TH) immunofluorescence). Scale bars, 500 μm and 20 μm. (b). Cell-attached recording of spontaneous activity in mCherry^+^ VTA neuron before, during, and after bath application of 1 µM clozapine-N-oxide (CNO). Example traces (left) and summary data (right). n = 4 cells from 3 mice, repeated measures one-way ANOVA, F (2, 6) = 9.634, p = 0.0134, Sidak’s multiple comparison test, Baseline vs CNO, p = 0.0104, CNO vs Washout, p = 0.0488. (c). Schematic of experimental timeline and pharmacological interventions in strong LH paradigm. (d). Left, summary data showing the percentage of failures to escape an escapable aversive shock across learning and treatment conditions for hM4Di AAV expressing DAT^iCre^ positive and negative littermates. Middle, within subject summary data for behavioral responses after ketamine treatment only, compared with ketamine + CNO, 4 hrs after treatment. Right, same but for 24 hrs after treatment. Two-way ANOVA, Sidak’s multiple comparison test, KET + CNO 4 hrs, p < 0.0001, KET + CNO 24 hrs, p = 0.0107, KET only, following 4, 24, and 72 hrs, p > 0.9, n = 7 - 8 animals. (e). Total distance traveled per minute in an open field locomotion assay for hM4Di^+^ animals before and after CNO treatment. Two-tailed paired t-test, p = 0.1473, n = 5 animals. (f). Summary data showing the percentage of failures to escape an escapable aversive shock in the baseline condition, for hM4Di-expressing mice before and after CNO treatment. n = 5 - 6 animals, Two-way ANOVA, Sidak’s multiple comparison test, Baseline, p = 0.9538, LH, p = 0.9947. (g). Left, schematic of experimental timeline and pharmacological interventions in weak LH paradigm. Right, summary data showing the percentage of failures to escape an escapable aversive shock across 6 days for two groups (white bar, hM4Di-controls; purple, hM4Di+). Two-way ANOVA, Sidak’s multiple comparison test, hM4Di+ vs hM4Di-, D1-D4, p > 0.5, D5, p < 0.0001, D6, p = 0.0001, n = 8 animals. *p < 0.05, *** p < 0.001, **** p < 0.0001. Error bars reflect SEM.

Mice conditionally expressing hM4Di in VTA DA neurons were compared to their Cre^-^ littermate controls in the strong LH paradigm. Here, two ketamine treatments were administered on sequential days – the first one along with CNO, and the second one without (Figure 3c). The expression of hM4Di in DA neurons did not change baseline behavior or learned helplessness induction (Figure 3d). However, inhibiting DA neurons by CNO application (3 mg/kg, b.w., i.p.) co-administered with ketamine (10 mg/kg, b.w., i.p.) blocked ketamine’s rescue of escape behaviors. This effect was evident 4 hrs after CNO/Ketamine treatment and persisted through 24 hrs. Yet, when ketamine was administered alone on the following day, the behavioral rescue in hM4Di^+^ mice was successful. Persistent LH was evident in multiple animals of both groups 72 hrs following the ketamine only treatment (last two bars, Figure 3d). Comparing behavioral responses following ketamine alone versus ketamine plus CNO, within animals at two separate time-points, showed that the responses of hM4Di^-^ animals lie around the unity line (Figure 3d). In contrast, hM4Di^+^ responses were above the unity line, reflecting selective efficacy of ketamine treatment in the absence of CNO. Locomotor behavior in hM4Di-expressing animals was not grossly altered by a single CNO administration, suggesting that the observed changes in escape actions are not due to reduced movement (Figure 3e).

Although chemogenetic inhibition of DA activity blocked the behavioral effect of ketamine after aversive learning, the administration of CNO in the absence of ketamine in separate groups of hM4Di^+^ mice did not alter the proportion of failures to escape in the baseline condition or after sLH (Figure 3f). These data support the alternative hypothesis that the effect of ketamine after LH specifically requires the associated restoration of DA signals. Although VTA DA activity varies with behavioral outcomes before aversive learning, innate escape actions may not fundamentally require DA activity. CNO co-administration likewise blocked the behavioral effects of ketamine in the relatively weaker learning paradigm (wLH, Figure 3g). These data support the necessity of VTA DA signaling for the behavioral reversal of aversive learning by ketamine, rather than for native escape responses in naïve animals.

## The function of mPFC DA release in aversive learning

The activity of DA terminals in mPFC is implicated in the processing of emotionally valenced information, including aversive stimuli^26,32,51^. In particular, the mPFC is linked to behavioral flexibility in humans and other animals^52,53^. Dopamine release in mPFC modulates its activity and plasticity^54^, altered in animal models of stress^15,35^. Emerging data suggest that mPFC is one of the key regions that mediate rapid antidepressant actions of ketamine^15,35,55^. To specifically evaluate DA signaling in mPFC, rather than Ca^2+^ influx, we turned to the recently developed genetically encoded DA sensor dLight1.1^56^. We created a Cre-dependent AAV expressing dLight1.1 under the control of the CBA promoter. This specificity was important, because different DA receptors are expressed in distinct subsets of mPFC neurons^57–59^. We evaluated dLight1.1 reported DA signal in mPFC Drd1^+^ neurons in Drd1^Cre^(FK150) mice that were virally transduced with AAV1.CBA.DIO.dLight1.1 and implanted with optical fibers (Figure 4a, b). We found robust DA transients in mPFC Drd1a^+^ neurons in response to foot shocks. The responses were blunted at the end of LH induction, and recovered with ketamine administration along with escape behavior (Figure 4c, d). EGFP-expressing controls showed no responses (Figure 4d). The timing of positive deflection peaks for dLight preceded that of VTA DA Ca^2+^ transients (Figure 1d). One potential explanation for this observation is that functionally significant amounts of DA could release from local axons prior to sufficient accumulation of calcium required for GCaMP6f readout. Altogether, our data suggest that dopamine signaling in mPFC, like somatic Ca^2+^ photometry, carries a signature of the animal’s behavioral state.

**Figure 4.**
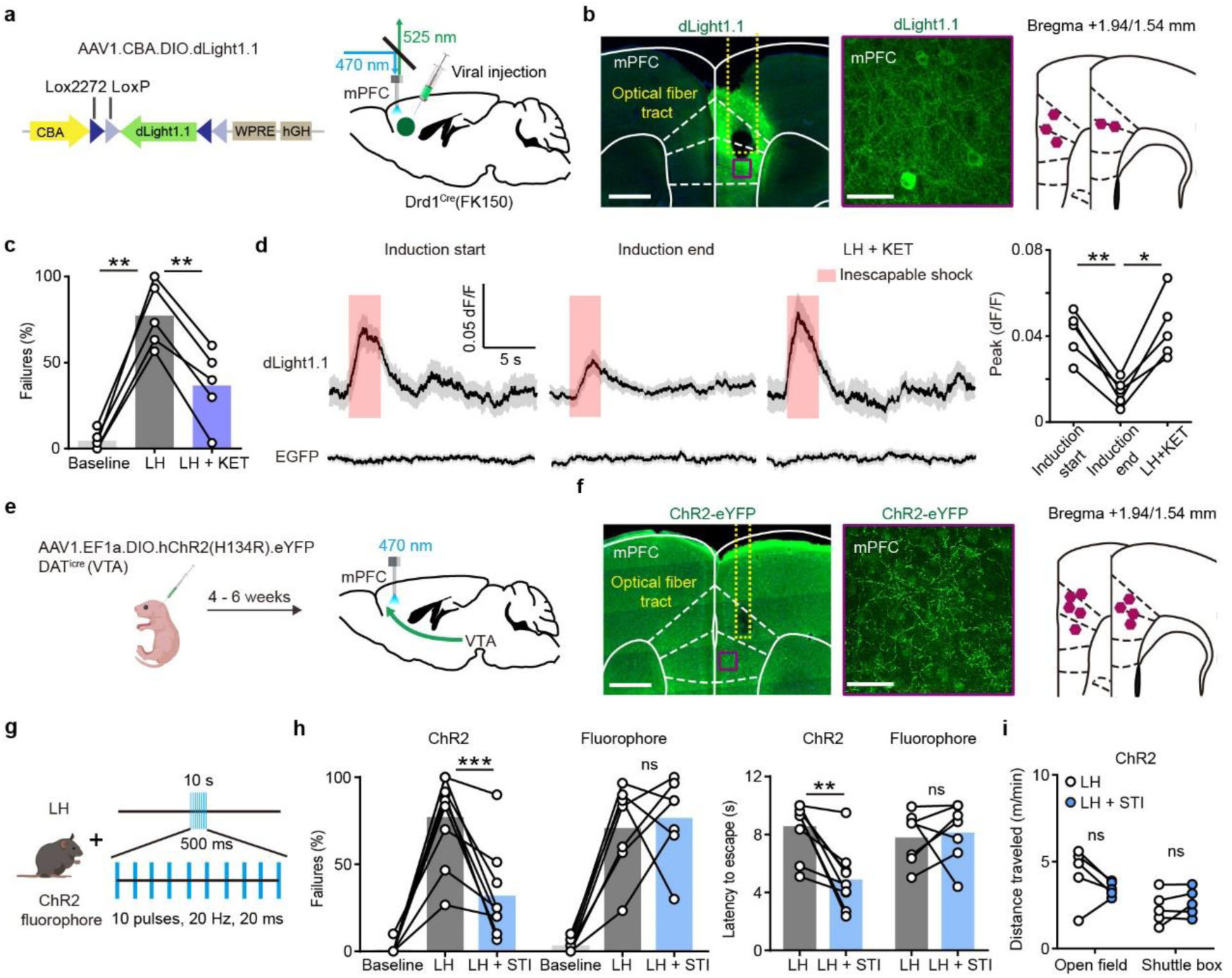
mPFC DA release dynamics and function in aversive learning. (a). AAV1.CBA.DIO.dLight1.1 construct and the schematic for viral transduction in the mPFC with subsequent fiber implant in Drd1^Cre^(FK150) mice. (b). Left, fiber placement illustration on a coronal section through mPFC, with a close up image of dLight1.1^+^ neurons and terminals (white dashed lines, Paxinos atlas overlay; yellow dashed lines, fiber track). Green, immunoenhanced dLight1.1; blue, Hoechst nucleic stain. Scale bars: 500 μm and 50 μm. Right, atlas location of fiber placement for each subject. (c). Summary data showing the percentage of failures to escape an escapable aversive shock in dLight1.1-expressing mice across phases of learning (Baseline, LH, and LH + KET). n = 5 animals, repeated measures one-way ANOVA, F (1.434, 5.735) = 51.75, p = 0.0003, Sidak’s multiple comparison test, Baseline vs LH, p = 0.0010, LH vs LH + KET, p = 0.0023. (d). Left top, average dLight1.1 transients in mPFC in one mouse, aligned to shock start time across learning phases (20 trials/condition, mean ± SEM; shock length, 3 sec, pink shading). Left bottom, EGFP controls. Right, within subject group data. Repeated measures one-way ANOVA, F (1.693, 6.773) = 24.16, p = 0.001, Sidak’s multiple comparison test, Baseline vs LH, p = 0.0036, LH vs LH + KET, p = 0.0107, n = 5 animals. (e). Schematic for viral transduction with Cre-dependent ChR2 AAV in the VTA and subsequent optogenetic fiber implant in mPFC. (f). Left, fiber placement illustration on a coronal section through mPFC, with a close up image of ChR2.eYFP terminals (white dashed lines, Paxinos atlas overlay; yellow dashed lines, fiber track). Green, immunoenhanced ChR2.eYFP; blue, Hoechst nucleic stain. Scale bars: 500 μm and 50 μm. Right, atlas location of fiber placement for each subject. (g). Schematic illustrating open loop optogenetic stimulation parameters (STI denotes optogenetic stimulation). (h). Left, summary data showing the percentage of failures to escape an escapable aversive shock in ChR2-expressing mice (n = 9) and fluorophore-expressing controls (n = 7) across phases of learning, Baseline, LH, and LH + STI. Right, summary data for latency to escape in LH compared with LH + STI conditions. Repeated measures two-way ANOVA, Sidak’s multiple comparison test, LH vs LH + STI, ChR2, p = 0.0002, Fluorophore, p = 0.9358. Latency to escape, LH vs LH + STI, ChR2, p = 0.0014, Fluorophore, p = 0.9248. (i). Left, locomotion in the open field and shuttle box (m/min) after learning with and without optogenetic stimulation. Repeated measures two-way ANOVA, Sidak’s multiple comparison test, open field, p = 0.1742, shuttle box, p = 0.7503, n = 5 mice. *p < 0.05, ** p < 0.01, *** p < 0.001. Error bars reflect SEM.

If the impairment of escape behavior after aversive learning is caused by blunted DA release in mPFC, increasing local DA release after learned helplessness should rescue escapes. To test this prediction, we optogenetically activated DA terminals in mPFC in animals with ChR2 expression restricted to VTA DA neurons. DAT^icre^ neonates were transduced with AAV1.EF1α.DIO.hChR2(H134R).eYFP or a control fluorophore and implanted with optical fibers 4-6 weeks after transduction (Figure 4e, f). Following learned helplessness induction, during the test session animals received a series of burst optogenetic stimuli at 20 Hz every 10 sec (10 pulses, 20 ms width, 500 ms duration), in the absence of ketamine treatment (Figure 4g). Note that the stimulation bursts were not timed relative to shocks and took place on either side of the shuttle box, decreasing the likelihood of forming conditioned place aversion. Optogenetic activation of DA axon terminals in mPFC significantly decreased the percentage of failures after LH, as well as latencies to escape (Figure 4h). The magnitude of these effects of optogenetic stimulation was comparable to ketamine (ChR2, 45% vs KET, 55%, percentage decrease in failures after LH). As previously observed for hM4Di-mediated inhibition of VTA DA neuron activity, optogenetic stimulation did not alter locomotion behavior in either the open field or the shuttle box (Figure 4i). Thus, enhancing DA release in mPFC, likely by recovering DA tone, is sufficient to rescue escape behavior after aversive learning.

## Evoked structural plasticity in mPFC is regulated by ketamine in a DA-dependent manner

Ketamine is widely believed to generate its therapeutic effects, at least in part, by modulating plasticity of mPFC neurons^5,11,12,17 60^. Several prior studies demonstrate that *in vivo* administration of ketamine enhances dendritic spine density^15,61–63^ and restores dendritic spine loss^18^. Whether ketamine directly increases spinogenesis and synaptogenesis, or alternatively decreases neuronal synapse maintenance and turnover remains unclear, although one recent study reported increased genesis with no changes in the elimination of dendritic spines, when ketamine was administered following chronic corticosterone treatment^18^. To determine the precise mechanism of ketamine regulation of structural plasticity and to evaluate whether changes in DA signaling contribute to ketamine effects in mPFC, we turned to focal induction of *de novo* spinogenesis in acute brain slices using 2-photon uncaging of glutamate. This spatiotemporally precise and high throughput plasticity assay is compatible with pharmacological and genetic manipulations, while maintaining a subset of circuit-level connections that would be impossible in a reduced culture system.

Acute slices of mPFC were prepared from P25-40 mice of both sexes following neonatal transduction of sparse EGFP expression accomplished by a combination of AAV1.hSyn.Cre and AAV8.FLEX.EGFP. We imaged EGFP-labeled dendrites of layer 5 pyramidal neurons in mPFC using 2-photon laser scanning microscopy (2PLSM, 910 nm). A second laser was tuned to 725 nm to locally uncage MNI-glutamate near dendrites to induce new dendritic spines (Figure 5a), as previously described for developing neurons in the striatum and superficial layers of sensory and motor cortex^50,64,65^. We carried out evoked spinogenesis assays in different mice at several time points (2-72 hrs) after a single dose of ketamine (10 mg/kg, i.p.) (Figure 5a).

**Figure 5.**
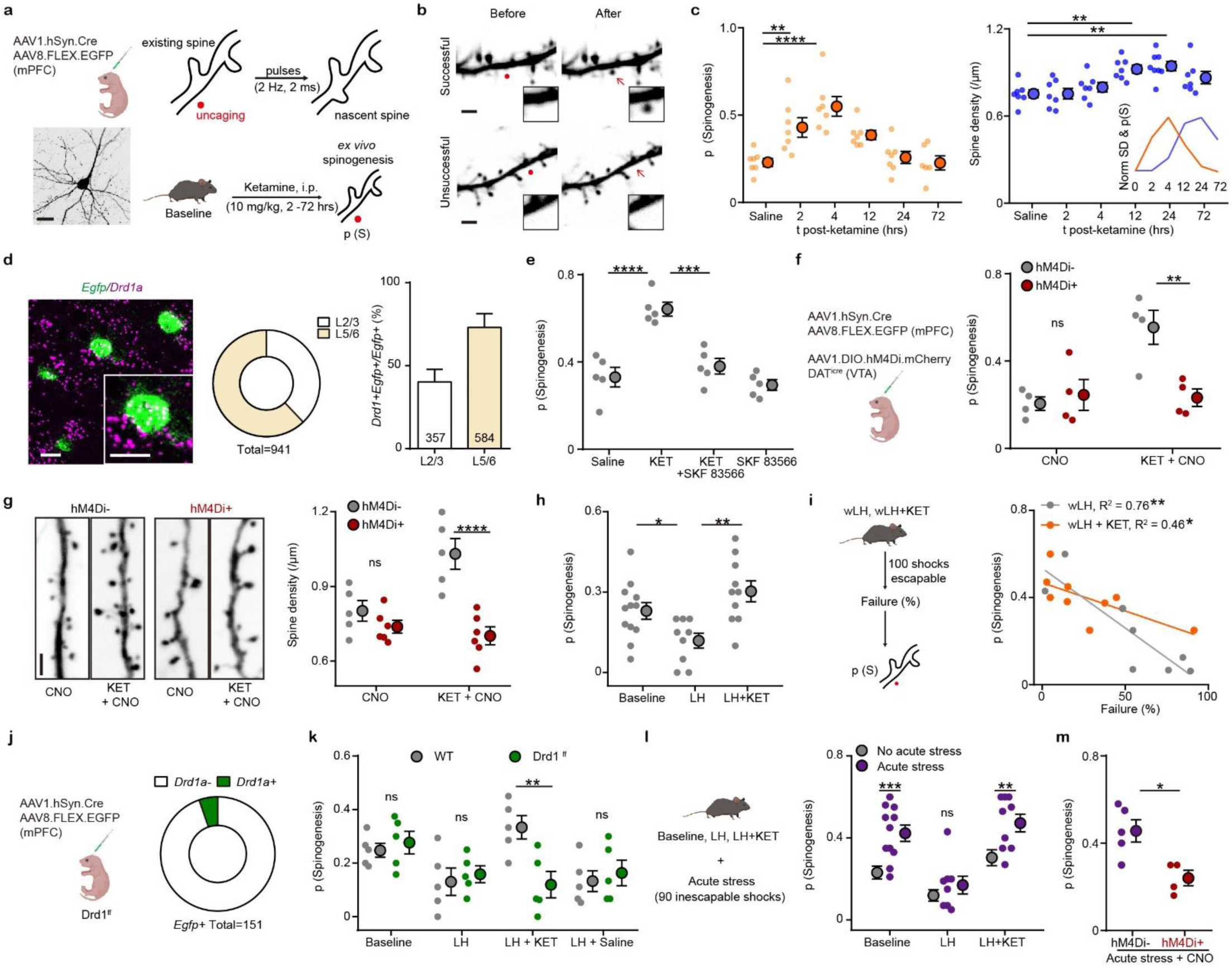
Ketamine regulates mPFC plasticity through a DA-dependent mechanism. (a). Schematic illustrating glutamate-evoked *de novo* spinogenesis platform. Left, viral transduction and an example EGFP^+^ pyramidal neuron in mPFC. Right, MNI-glutamate uncaging parameters for the induction of new dendritic spines and schematic illustrating pharmacological manipulation. Scale bar, 50 µm. (b). Example 2PLSM images of successful and unsuccessful induction trials of *de novo* spinogenesis. Red circles, uncaging sites. Black rectangle, close up images of local dendritic segments before and after glutamate uncaging. Scale bar, 2 µm. (c). Left, timecourse of evoked spinogenesis probability on deep layer mPFC neurons in mice treated with either saline or ketamine (i.p. 10 mg/kg, acute slice preparation 2-72 hrs after treatment). Each small circle, aggregate probability of evoked spinogenesis from a single animal. Large circle, group data. n = 6 - 7 animals/time point, 15 - 25 trials/animal, one-way ANOVA, F (5, 35) = 9.895, p < 0.0001, Sidak’s multiple comparison test vs Saline, 2 hrs p = 0.076, 4 hrs, p < 0.0001, 12 hrs, p = 0.0532, 24/72 hrs, p > 0.9. Right, same as left but for dendritic spine density. n = 7 - 8 animals/time point, one-way ANOVA, F (5, 37) = 6.319, p = 0.0002, Sidak’s multiple comparison test vs Saline, 2/4 hrs p > 0.8, 12 hrs, p = 0.0056, 24 hrs, p = 0.0011, 72 hrs, p = 0.1271. Inset, normalized time course of changes in evoked spinogenesis (orange) and dendritic spine density (blue). (d). Left, fluorescence *in situ* hybridization (FISH) image showing *Drd1a* mRNA in *Egfp* mRNA expressing mPFC cells. Inset, close up of a single neuron. Middle, the proportional distribution of *Egfp* mRNA expression across superficial and deep cortical layers. Right, quantification of the percentage of *Drd1a*^*+*^ cells among *Egfp*^*+*^ cells in superficial and deep cortical layers. n = 4 animals; *Drd1*^*+*^*Egfp*^*+*^*/Egfp*^*+*^: Layer 2/3, 40.2% ± 7.5%, layer 5/6, 73.1% ± 8.3%. Scale bar, 20 µm. (e). Probability of glutamate-evoked spinogenesis on deep layer mPFC neurons in mice treated with Saline, KET (10 mg/kg), KET + SKF 83566 (10 mg/kg), or SKF 83566 alone. Each small circle, aggregate probability of evoked spinogenesis from a single animal. Large circle, group data. One-way ANOVA, p < 0.0001, F (3, 16) = 20.29, Sidak’s multiple comparison test, Saline vs KET, p < 0.0001, KET vs KET + SKF83566, p = 0.0002, Saline vs SKF83566, p = 0.8574. (f). Left, schematic illustrating triple viral transduction strategy for evoked spinogenesis with DA neuron inhibition. Right, probability of spinogenesis on deep layer mPFC neurons in DAT^iCre+^ and DAT^iCre-^ animals treated with CNO (3 mg/kg) across conditions (baseline, KET). n = 4 animals/condition as shown in plots, two-way ANOVA, Sidak’s multiple comparison test, Cre-vs Cre+, CNO, p = 0.8686, CNO + KET, p = 0.0042. (g). Left, example confocal images of EGFP expression in dendrites of deep layer mPFC pyramidal neurons, in response to CNO and ketamine treatment, as noted. Scale, 2 µm. Right, same as (f) but for dendritic spine density. n = 5 - 6 animals/condition as shown in plots, two-way ANOVA, Sidak’s multiple comparison test, Cre^-^ vs Cre^+^, CNO, p = 0.5005, CNO + KET, p < 0.0001. Scale bar, 2 µm. (h). Probability of glutamate-evoked spinogenesis on deep layer mPFC neurons in distinct stages of aversive learning (baseline, LH, LH + KET). n = 9 - 12 animals/condition as shown in plots, one-way ANOVA, F (2, 28) = 7.146, p = 0.0031, Sidak’s multiple comparison test, Baseline vs LH, p = 0.0496, LH vs LH + KET, p = 0.0016. (i). Left, schematic illustrating glutamate-evoked spinogenesis assay after wLH and wLH + KET in separate groups of mice. Right, correlation between failures to escape and the probability of spinogenesis after learning and following ketamine treatment. wLH, R^2^ = 0.76, p = 0.0051, n = 8 animals. wLH + KET, R^2^ = 0.46, p = 0.0439, n = 9 animals. (j). Left, schematic illustrating dual viral transduction strategy with sparse genetic manipulation of Drd1 receptor expression in Drd1^ff^ mice. Right, quantification of the percentage of *Drd1a*^*+*^ cells among *Egfp*^*+*^ cells in mPFC. 5% *Drd1a*^*+*^ and 95% *Drd1a*^-^ among 151 Egfp^+^ cells from 2 animals. (k). Probability of glutamate-evoked spinogenesis on deep layer mPFC neurons in distinct stages of aversive learning (baseline, LH, LH + KET, LH + saline) in wild type and Drd1^ff^ mice. Two-way ANOVA, Sidak’s multiple comparison test, WT vs Drd1^ff^, LH + KET, p = 0.0043, Baseline, LH and LH + Saline, p > 0.9, n = 5 animals. (l). Left, schematic illustrating acute stress manipulation on the background of aversive learning states (Baseline, LH, LH+KET). n = 8 - 12 animals/condition as shown in plots, two-way ANOVA, Sidak’s multiple comparison test, Baseline vs Baseline stressed, p = 0.0007. LH vs LH stressed, p = 0.7548. LH +KET vs LH + KET stressed, p = 0.0083. Grey dots, summary data from panel h. (m). Probability of glutamate-evoked spinogenesis in hM4Di- and hM4Di+ mice following acute stress and CNO treatment. Unpaired two-tailed t test, p = 0.0132, n = 4 - 5 animals. *p < 0.05, ** p < 0.01, *** p < 0.001 **** p < 0.0001. Error bars reflect SEM.

Successful and unsuccessful induction trials of *de novo* spinogenesis were distinguished in z-stack projections through a dendritic segment prior to and following the brief induction protocol (< 30 sec) of up to 40 uncaging pulses (Figure 5b). In order to be classified as newly induced dendritic spines, the new membrane protrusions had to satisfy several criteria based on location and fluorescence intensity, relative to parent dendrite and pre-existing dendritic spines (methods and Extended Data Figure 3a-c). A single *in vivo* administration of ketamine in naïve animals enhanced evoked *de novo* spinogenesis 2 and 4 hours after treatment (Figure 5c). This effect was transient – by the 12 hr time point after ketamine, the probability of spinogenesis decreased back to baseline levels. In addition, dendritic spine density was quantified at the same time points. In contrast to the rapid, transient changes in evoked spinogenesis, the increase in spine density was delayed until 12 hrs after treatment (Figure 5c). Because the increase in glutamate-evoked spinogenesis precedes the increase in dendritic spine density, the accumulation of changes in the plasticity potential is a likely contributing factor to post-ketamine changes in dendritic spine density^13,17,18,61^.

To determine whether ketamine’s effect on evoked plasticity is mediated by the activation of DA receptors, we first verified the expression of Drd1a receptors in EGFP-expressing neurons. Consistent with prior reports^66^, the majority of pyramidal neurons in the deep layers of mPFC express *Drd1a* mRNA (Figure 5d). We compared glutamate-evoked spinogenesis after administering ketamine alone, or in conjunction with a Drd1 receptor antagonist SKF 83566 (10 mg/kg i.p., 2 hrs prior to *ex vivo* experiments). We found that antagonizing Drd1 receptors blocked ketamine’s potentiation of evoked spinogenesis, while the antagonist treatment alone had no effect relative to baseline (Figure 5e). Thus, while the activation of Drd1 receptors in this neuronal population is not required for baseline glutamate-evoked plasticity, it appears to be necessary for ketamine’s enhancement of evoked spinogenesis. As a complement to pharmacological manipulation of Drd1 receptors, since SKF83566 treatment *in vivo* broadly alters Drd1 activation and locomotor behavior, we turned again to chemogenetic inhibition of VTA DA neurons to modulate downstream DA release. Inhibiting hM4Di^+^ VTA DA neurons with CNO (3 mg/kg, i.p.) blocked spinogenesis-enhancing effects of ketamine (Figure 5f). Yet, as for the pharmacological Drd1 receptor blockade *in vivo*, we observed no effects of CNO treatment on evoked spinogenesis in the absence of ketamine. These observations are consistent with a model where the genesis of new dendritic spines and synapses mechanistically depends on glutamate, but the enhancement of this plasticity requires the activation of protein kinase A (PKA) via Gα_s_-coupled receptors, as has been previously reported for the striatum during early postnatal development^50^. In addition to blocking ketamine-mediated enhancement of evoked spinogenesis, transient inhibition of VTA DA neuron activity (a single CNO dose + ketamine) also abolished the delayed increase of spine density 24 hrs after ketamine (Figure 5g). These data show that in the absence of behavioral manipulations, Drd1 activation and VTA DA activity regulate changes in spinogenesis and dendritic spine density, mediating the effects of ketamine on plasticity in mPFC.

In order to connect ketamine’s effects on evoked plasticity to aversive learning, we next tested glutamate-evoked spinogenesis in baseline, LH, and LH + KET conditions using the strong LH paradigm. The probability of glutamate evoked spinogenesis decreased relative to baseline in sLH trained mice, while ketamine treatment restored baseline potential for plasticity (Figure 5h). To correlate individual-level behavioral outcomes with evoked plasticity, we performed *de novo* spinogenesis assays in wLH trained animals with or without subsequent ketamine treatment. We found that the probability of evoked spinogenesis negatively correlated with the percentage of failure to escape in both conditions, providing a more direct link between spinogenesis and behavior (Figure 5i). To specifically manipulate Drd1 receptor expression in mPFC without affecting the global DA system, we conditionally knocked out Drd1 receptor by co-expressing Cre recombinase and floxed EGFP in mice with Drd1-floxed genetic background (Figure 5j). We validated the conditional knock-out by verifying the expression of *Drd1a* mRNA in EGFP-expressing neurons (Figure 5j and Extended Data Figure 3d). Sparse genetic depletion of Drd1 receptor in mPFC pyramidal neurons abolished the effect on spinogenesis induced by ketamine in sLH trained animals, without changing the probability of spinogenesis in mice in the baseline or sLH conditions (Figure 5k).

We then sought a deeper understanding of the mechanistic relationship between ketamine effects on plasticity and experience-dependent changes in dopaminergic system, where initial unpleasant stimuli elicit robust responses, but fail to do so after aversive learning. We examined glutamate evoked *de novo* spinogenesis on mPFC neurons from mice across distinct behavioral states of aversive learning, with or without additional acute stress. The inclusion of acute stress on the background of different aversive learning states was motivated by two factors. First, the acute stress of the initial shock stimuli in the baseline elicits robust DA responses, which are blunted after LH. Given that the observed increases in glutamate-evoked spinogenesis require DA signaling via Drd1 receptor, we hypothesize that acute stress should potentiate spinogenesis in the baseline condition, but not after LH. Second, prior studies of experience-induced structural changes in the brain report distinct and sometimes opposite effects of acute and prolonged stress^16^. We chose to use the same type of foot shock stimuli for the acute stress as for the aversive learning induction procedure, in order to specifically determine the effects of prior experience on plasticity. The first exposure to acute stress of 90 foot shocks (3 sec, 0.3 mA, ∼15 min total) significantly enhanced evoked *de novo* spinogenesis on pyramidal neurons in mPFC. This observation is consistent with our data on DA mediation of enhanced mPFC plasticity. DA transients are blunted after LH, raising the question of whether this blunting is sufficient to block the effect of acute stress on plasticity. Indeed, following aversive learning induced by prolonged exposure to inescapable foot shocks (2 days, 360 shocks/day, ∼140 min total), the plasticity-promoting effect of additional acute stress of 90 foot shocks was abolished. Finally, ketamine rescued this potentiation of evoked plasticity by acute stress (Figure 5l). These effects were similar in males and females, depicted in binary grid plots of individual induction trials (Extended Data Figure 3e). We confirmed that the effect of acute stress on evoked plasticity is DA-dependent by using chemogenetic inhibition of VTA DA neurons (Figure 5m).

The final series of plasticity experiments addressed the mechanism by which DA enhances glutamate-evoked spinogenesis. Drd1 receptor activation is known to regulate glutamatergic synapse and dendritic spine formation in the developing striatum through PKA^50^. Analogously, we found that bath application of Drd1 agonist SKF 81297 (1 μM) promotes glutamate-evoked spinogenesis on mPFC pyramidal neurons (Extended Data Figure 4a, b). This effect required Drd1a signaling, since Drd1a cKO abolished the enhancement of spinogenesis. Suppression of PKA activity by either bath application of H-89 (10 μM) or over-expression of endogenous PKA inhibitor (PKIα) in mPFC pyramidal neurons blocked changes in spinogenesis induced by SKF 81297 (Extended Data Figure 4b). In addition, *in vivo* pre-treatment with ketamine (10 mg/kg, i.p.) occluded the enhancement of spinogenesis by SKF 81297 (Extended Data Figure 4c). Furthermore, the plasticity-promoting effect of ketamine was blocked by over-expression of PKIα (Extended Data Figure 4d, e). The link between Drd1a-PKA signaling and dendritic spine regulation is largely contributed by PKA downstream targets including GTPases and their regulatory proteins involved in cytoskeleton remodeling^67^ (Extended Data Figure 4f). Altogether, our results reveal that aversive experience and ketamine modulate mPFC plasticity through mechanisms that require DA, Drd1a signaling, and PKA activity.

## Closed loop interactions between VTA and mPFC in aversive learning

While we find that systemically inhibiting VTA DA neuron activity abolishes the behavioral effect of ketamine and mPFC plasticity changes, whether the behavioral response to ketamine primarily operates through local dopamine release in mPFC remains unclear. To achieve local inhibition of dopamine release, we infused CNO into the mPFC of mice expressing hM4Di in VTA DA neurons and their terminals in mPFC, to reduce axonal release probability^47,68,69^. DAT^iCre^ neonates were transduced with AAV1.CBA.DIO.hM4Di.mCherry in the VTA, and cannulae were implanted bilaterally over mPFC (at ∼P50) in order to locally deliver 1 mM CNO (1 µl for each side) (Figure 6a and Extended Data Figure 5a). A high density of hM4Di mCherry expression in mPFC terminals was observed in immunoenhanced fixed tissue sections (Figure 6b). Local infusion of CNO in mPFC blocked the behavioral effect of ketamine (10 mg/kg, i.p.) in the aversive learning paradigm, while ketamine alone was sufficient to rescue escape behavior (Figure 6c and Extended Data Figure 5b). These data are consistent with our prior experiments on the inhibition of VTA DA neuronal excitability (Figure 3, i.p. injection of CNO), but the sufficiency of an mPFC-specific activation of hM4Di confirms mPFC as the key site of control over behavioral changes driven by ketamine.

**Figure 6.**
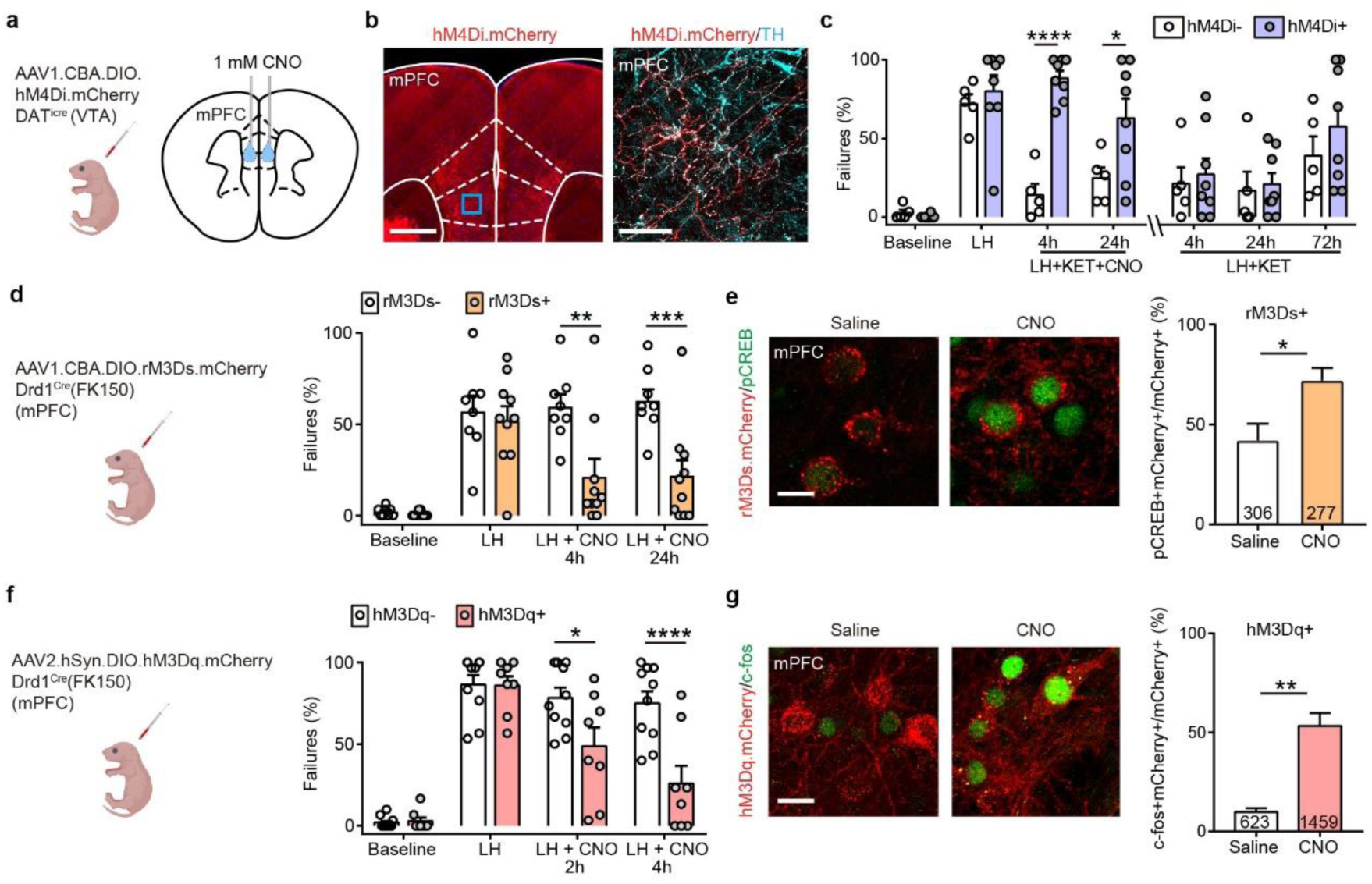
Activity of local DA terminals and Drd1^+^ neurons in mPFC mediates behavioral responses to ketamine. (a). Left, schematic illustrating viral transduction strategy. Right, local CNO infusion in mPFC (1 mM, 1 μl). (b). Left, immunoenhanced image of hM4Di.mCherry^+^ DAT^+^ terminals in mPFC. Right, mCherry^+^ terminals colocalize with a subset of tyrosine hydroxylase (TH) expressing axons. Scale bars, 500 μm and 50 μm. (c). Summary data showing the percentage of failures to escape an escapable aversive shock across learning and treatment conditions for hM4Di-expressing DAT^iCre^ positive and negative littermates. n = 5 animals for Cre^-^, 8 animals for Cre^+^, two-way ANOVA, Sidak’s multiple comparison test. KET + CNO 4 hrs, p < 0.0001, KET + CNO 24 hrs, p = 0.0476, KET + only 4, 24, and 72 hrs, p > 0.7. (d). Left, schematic illustrating viral transduction strategy. Right, Summary data showing the percentage of failures to escape an escapable aversive shock in Drd1-Cre^+^ and Drd1-Cre^-^ mice expressing rM3Ds across phases of learning and after CNO treatment (Baseline, LH, LH + CNO 4 hrs, and LH + CNO 24 hrs). n = 8 - 10 animals/condition, two-way ANOVA, Sidak’s multiple comparison test, Cre^+^ vs Cre^-^, LH + CNO 4 hrs, p = 0.0018, LH + CNO 24 hrs, p = 0.0007, Baseline/LH, p > 0.9. (e). Left, colocalization of pCREB immunolabeling and rM3Ds.mCherry expression in mPFC after Saline/CNO treatment in Drd1 Cre^+^ mice. Right, the quantification of percentage of pCREB^*+*^ cells among mCherry^*+*^ cells. Scale bar, 20 μm. n = 3 animals/condition cell number as noted in each bar, two-tailed unpaired t-test, p = 0.0455. (f). Left, schematic illustrating viral transduction strategy. Right, summary data showing the percentage of failures to escape an escapable aversive shock in Drd1-Cre^+^ and Drd1-Cre^-^ mice expressing hM3Dq across phases of learning and after CNO treatment (Baseline, LH, LH + CNO 2 hrs, and LH + CNO 4 hrs). n = 8 animals for Cre^-^, n = 10 mice for Cre^+^, two-way ANOVA, Sidak’s multiple comparison test, LH + CNO 2 hrs, p = 0.0157, LH + CNO 4 hrs, p <0.0001, Baseline/LH, p > 0.9. (g). Left, colocalization of c-fos immunolabeling in hM3Dq.mCherry^+^ mPFC neurons in saline and CNO treated Drd1 Cre^+^, hM3Dq-expresing mice. Right, the quantification of percentage of c-fos^*+*^ cells among mCherry^*+*^ cells. Scale bar, 20 μm. n = 3 mice/condition, two-tailed unpaired t-test, p = 0.0031. *p < 0.05, ** p < 0.01, *** p < 0.001, **** p < 0.0001. Error bars reflect SEM.

The activation of Drd1 receptors initiates Gα_s_ mediated PKA signaling cascades, which enhance spinogenesis, synaptic transmission, and neuronal activity^49,50,66,70^. We therefore tested whether selective activation of Gα_s_ signaling in mPFC Drd1 expressing neurons could rescue escape behavior after aversive learning. We relied on the Gα_s_-coupled rM3D DREADD, expressing AAV1.CBA.DIO.rM3Ds.mCherry in Drd1-Cre-FK150 mice (Figure 6d). The expression of rM3Ds alone did not change baseline escape and failure rates, or the magnitude of aversive learning. After LH induction, a single i.p. dose of CNO was sufficient to rescue escape behavior 4 hours after treatment, lasting at least 24 hours (Figure 6d). Activating Gα_s_ signaling in Drd1a expressing neurons *in vivo* significantly increased phosphorylation of CREB, which is typically induced by Gα_s_-coupled cascades (Figure 6e). Since the activation of other G protein-coupled cascades can also lead to an increase in cellular excitability, in a complementary experiment we used the Gα_q_-coupled hM3D to directly enhance the excitability of Drd1 expressing neurons in mPFC following aversive learning (Figure 6f). The expression and functional activation of hM3Dq were validated by immunohistochemistry and electrophysiology (Extended Data Figure 5c, d). The expression of hM3Dq also did not change baseline escape/failure rates, or the magnitude of aversive learning. After LH induction, a single i.p. dose of CNO was sufficient to rescue escape behavior within 2 hours, with a larger effect observed 4 hours after CNO treatment (Figure 6f). The activation of hM3Dq in Drd1-positive mPFC neurons did not alter locomotion (Extended Data Figure 5e). At the end of the experiment, the successful enhancement of neuronal activity was further validated by evaluating immediate early gene product c-fos expression in mCherry^+^ neurons (Figure 6g). In addition to our results, a recently published study showed that optogenetic activation of Drd1^+^ mPFC neurons decreases immobility time in the forced swim test, suggesting the possibility that these Drd1-expressing neurons may broadly regulate aversive or active coping responses^55^. Altogether, our data demonstrate that activating DA-sensing mPFC neurons mimics the behavioral and plasticity-promoting effects of ketamine.

While mPFC receives dopaminergic input from the VTA, it also sends excitatory glutamatergic projections back to VTA DA neurons, based on trans-synaptic tracing analyses and optogenetic assays^51,71–73^. Whether this recurrent VTA-mPFC circuit reflects a closed-loop interaction, involving the same neurons in the VTA and Drd1^+^ neurons in mPFC, is not known. To label neurons that send projections to VTA, we injected red retrograde tracer fluorescent beads in the VTA (Extended Data Figure 6a, b). Combined retrograde tracing and FISH experiments demonstrate that the majority of retrobead^+^ neurons in layer V of mPFC express *Drd1a* mRNA, suggesting they receive dopaminergic inputs. In contrast, a small fraction of the retrobead^+^ neurons in layers 2/3 of mPFC have *Drd1a* mRNA (Extended Data Figure 6c). To determine whether mPFC projecting VTA neurons also receive mPFC projections, we expressed AAV8.FLEX.EGFP in Drd1^Cre^(FK150) animals and injected retrobeads in mPFC (∼P30) (Extended Data Figure 6d). Consistent with previous reports^73^, most but not all mPFC-projecting VTA neurons express tyrosine hydroxylase (TH), indicating their dopaminergic identity (Extended Data Figure 6e). Anatomical tracing results show that mPFC-targeting VTA neurons reside in a region of dense projections from mPFC Drd1^+^ neurons (Extended Data Figure 6f). Since glutamate co-transmission from VTA DA terminals has been reported in nucleus accumbens^74^, we recorded light-evoked, pharmacologically isolated, excitatory post-synaptic currents (EPSCs) in layer V mPFC pyramidal neurons of mice transduced with Cre-dependent ChR2 in VTA DA neurons (Extended Data Figure 6g). No time-locked or delayed light-evoked EPSCs were observed in the presence of a GABA(A) receptor antagonist in neurons held at -70 mV (Extended Data Figure 6h), arguing for selective release of DA from the VTA onto deep layer mPFC pyramidal neurons. These data support the existence of a recurrent closed loop between VTA DA and mPFC DA-sensing neurons.

Results from several recent studies support a model where ketamine disinhibits mPFC pyramidal neurons by suppressing the activity of local inhibitory interneurons^4,8,9,75,76^. To directly test this hypothesis in our paradigm for DA-sensing mPFC neurons, we used *in vivo* fiber photometry to monitor the activity of either GABAergic or Drd1^+^ neurons in mPFC before and after ketamine treatment. To restrict the expression of GCaMP6f, we transduced AAV1.CAG.FLEX.GCaMP6f in either VGAT^iCre^ or Drd1^Cre^(FK150) mice (Extended Data Figure 7a, b, e, f). Population activity of GABAergic neurons was rapidly reduced after ketamine treatment (Extended Data Figure 7c, d). Conversely, we observed an enhancement in Ca^2+^ transients in cortical Drd1^+^ neurons (Extended Data Figure 8g, h), consistent with the disinhibition model proposed for ketamine effects on cortical activity.

To confirm the mPFC as a key input for ketamine regulation of VTA DA neuron activity, we carried out two additional experiments. First, *ex vivo* application of ketamine (50 μM) in acute VTA brain slices did not alter the power or frequency tuning of spontaneous Ca^2+^ dynamics^77^ in DA neurons (Extended Data Figure 8a-f). Second, *in vivo* local infusion of ketamine into mPFC (12.5 µg, 500 nl) sufficed to recover Ca^2+^ activity signatures in VTA DA neurons after aversive learning, in parallel with behavioral rescue (Extended Data Figure 8g, h). Altogether, our results support a recurrent circuit model that provides a unifying framework for ketamine effects on neuromodulation and plasticity in regulating behavior after aversive learning (Extended Data Figure 8i).

## Discussion

Animals must learn to avoid dangerous stimuli in nature in order to survive, underscoring the ethological significance of aversive learning. However, overlearned responses can be maladaptive. Prolonged exposure to stressful events can lead to learned helplessness^25,39,78^, where an individual fails to avoid unpleasant stimuli. Here, we demonstrate that aversive learning restructures neuronal activity in the VTA DA system, associated with a shift in behavioral outcome distribution. A single low-dose ketamine treatment suffices to temporarily normalize both DA signaling and behavior, with the efficacy of treatment correlating with VTA DA activity features prior to learning. Bidirectional manipulations of somatic or axonal DA activity occlude or mimic the effects of ketamine. As evidenced by glutamate uncaging-evoked formation of new dendritic spines on deep layer pyramidal neurons, ketamine modulates mPFC synaptic structural plasticity in a DA-dependent manner. Underscoring the link between structural synaptic plasticity and behavior, the probability of glutamate-evoked spinogenesis on mPFC neurons correlated with individual escape behavior both after learning and following ketamine administration.

Multivalenced encoding of information by VTA DA neurons and their projections is now well established^26,27,79–81^. In our framework, the differences in VTA DA activity between success and failure outcomes map to the tendency to escape aversive stimuli. Accordingly, decreases in the differences between VTA DA activity on escape and failure trials after aversive learning are associated with increased failures to escape avoidable stressful stimuli. Building on these correlative observations, a series of bidirectional perturbations of activity in DA neurons and DA-sensing populations in mPFC confirms a causal relationship between learning, DA activity, and ketamine effects. Chemogenetic inhibition of VTA DA neuron activity results in increased failures to escape after ketamine treatment. Conversely, optogeneticaly evoked DA release in mPFC or chemogenetic activation of downstream Drd1 receptor-positive neurons suffice to mimic ketamine’s behavioral effects. Altogether, our model shows that DA activity changes in aversive learning paradigms, and controlling DA dynamics drives behavioral outcomes.

Prior data from our lab^72^ and others^73,82,83^ confirm that VTA DA neurons receive excitatory mPFC inputs. Glutamate release from mPFC terminals elicits excitatory responses in VTA DA neurons^84–87^, driving synchronized activity across VTA and mPFC, as well as other limbic structures^87^. VTA DA neurons receiving mPFC inputs appear to form a part of a recurrent circuit: they project to mPFC, based on evidence from electron microscopy reconstructions^82^ and consistent with our anatomical observations. Ketamine’s effects on neural activity and behavior must be understood in the context of this recurrent system. Results from our study and others^4,8,9,75,76^ provide direct evidence that ketamine administration induces rapid disinhibition of cortical pyramidal neurons. Ketamine enhances VTA DA activity not through direct effects on these neurons, but by altering mPFC activity. Other potential circuit mechanisms for ketamine’s enhancement of VTA DA activity include ventral hippocampal inputs to nucleus accumbens and lateral habenula projections to the VTA, which are modulated by stressful experience and ketamine^34,88,89^.

Recent evidence from human brain imaging studies suggests that major depressive disorder is associated with reduced synaptic connections, which are likely linked to changes in network functional connectivity^90^. The long-standing theory for how stressful experience alters mPFC dendritic spines and synapses is an inverse parabolic function. Structural plasticity is enhanced by acute stress, but impaired after prolonged or chronic aversive experiences^16^. These insights have been gained primarily from studies using correlative analyses of dendritic spine density, which do not distinguish changes in new spine genesis from maintenance and turnover. In our results, acute stress enhances the probability of glutamate-evoked spinogenesis, while the relatively prolonged stress of aversive learning dampens it, with ketamine recovering both the baseline spinogenesis capacity, as well as the potentiation by acute stress. This ketamine modulation of the rules for plasticity – or metaplasticity – depends on DA signaling. Although the effects of stress on neural systems are certainly varied, our data highlight the critical function of DA in experience-dependent mPFC structural plasticity.

Our observations of DA signaling mediation of dendritic spine plasticity after ketamine in mPFC may reflect broadly conserved mechanisms in the brain, where DA controls activity-induced plasticity of dendritic spines and excitatory synapse formation. Our prior data demonstrate that, during development, DA regulates the formation of dendritic spines and excitatory synapses in striatal direct pathway spiny projection neurons expressing Drd1 receptors^50^. The activation of Drd1 receptors stimulates Gα_s_ signaling cascades, increasing cAMP production and PKA activity. Analogously, DA promotes glutamate-evoked spinogenesis on mPFC pyramidal neurons through Drd1 receptor activation and changes in PKA activity. Given that actin dynamics are important for dendritic spine formation and shape regulation^91^, the mechanistic link between Drd1-PKA signaling and dendritic spine formation likely involves cytoskeleton remodeling proteins. Indeed, PKA modulates the activity of small GTPases (*e.g*., Rap1, Rac1, Cdc42, among others) known to regulate dendritic spines^67^ through guanine nucleotide exchange factors (GEFs) and GTPase-activating proteins (GAPs)^92,93^. Specific molecular effectors responsible for ketamine-induced changes in synaptic and dendritic spine plasticity remain to be elucidated and may provide new clinical targets.

Glutamate-evoked interrogation of plasticity on genetically targeted neurons offers unique strengths as a structural plasticity readout. Besides dissociating *de novo* genesis and elimination of dendritic spines and synapses, this assay facilitates pharmacological and genetic mechanism dissection and is compatible with correlative analyses of behavior. Our observations demonstrate a temporal antecedence of spinogenesis increase relative to changes in dendritic spine density, suggesting that the changes in spine density *in vivo* can be due to a prior, accumulating change in glutamatergic activity-dependent spinogenesis. Recent work demonstrates that newly formed dendritic spines are required to maintain the behavioral effect of ketamine after chronic corticosterone administration^18^, establishing a causal link between the increase in new spine formation and ketamine’s behavioral effects. These findings highlight the importance of new dendritic spine formation for behavior regulation. Exactly how new dendritic spines stabilize and contribute to behavior after ketamine treatment may reveal how ketamine’s effects last days beyond its bioavailability. Since our experiments were carried out in young animals and neural plasticity dynamics are known to change across age^94–96^, the efficacy of ketamine treatment could vary in clinical populations as a function of age, even if mechanisms of action are conserved.

Drugs that modulate the dopaminergic system represent first line therapies for a large number of diverse neurological and mental health conditions that are comorbid with major depressive disorder, including Parkinson’s disease, schizophrenia, OCD, ADHD, and eating disorders^97–101^. The recent FDA approval of esketamine for the treatment of major depressive disorder is a major expansion of its clinical use. Our results on dopaminergic mediation of ketamine’s effects on behavior and plasticity suggest the possibility of ketamine’s differential clinical effects in the large population of patients receiving exogenous DA precursors (e.g. L-DOPA), re-uptake inhibitors, as well as dopaminergic receptor agonists and antagonists.

## Acknowledgements

The authors are grateful to Lindsey Butler for mouse colony management, Northwestern Biological Imaging Facility and Dr. Tiffany Schmidt for confocal microscope access. This work was supported by the Rita Allen Foundation Scholar Award, NINDS R01NS107539, Searle Scholar Award, the Beckman Young Investigator Award, William and Bernice E. Bumpus Young Innovator Award, NARSAD Young Investigator, and P&S Fund Grant (all Y.K.). M.W was supported as an affiliate fellow of the NIH T32 AG20506, S.M. is a fellow of the National Science Foundation Graduate Research Fellowship DGE-1842165, and V.D. is a predoctoral fellow of the American Heart Association (19PRE34380056).

## Author Contributions

M.W. and Y.K. designed the study. M.W. carried out most experiments and analyses in the study. S.M. made major contributions to behavioral and photometry analyses, V.D. created new plasmids, viruses, and contributed to experimental analyses. P.H. and L.X contributed to morphological analyses and functional validation of DREADD tools. All authors participated in writing the manuscript.

## Methods

### Mouse strains and genotyping

Animals were handled according to protocols approved by the Northwestern University Animal Care and Use Committee. Weanling and young adult male and female mice (postnatal days 21-80) were used in this study. Approximately equal numbers of males and females were used for every experiment. All mice were group-housed, with standard feeding, light-dark cycle, and enrichment procedures; littermates were randomly assigned to conditions. C57BL/6 mice used for breeding and backcrossing were acquired from Charles River (Wilmington, MA), and all other mouse lines were acquired from the Jackson Laboratory (Bell Harbor, ME) and bred in house.

B6.SJL-Slc6a3^tm1.1(cre)Bkmn^/J mice, which express Cre recombinase under control of the dopamine transporter promoter, are referred to as DAT^iCre 102^; B6.FVB(Cg)-Tg(Drd1-cre)FK150Gsat/Mmucd mice, which express Cre recombinase under control of the dopamine Drd1a receptor promoter, are referred to as Drd1^Cre^(FK150)^103^; Drd1^tm2.1Stl^ floxed mutant mice that possess loxP sites flanking the single exon of the Drd1a gene, are referred to as Drd1^ff 104^; Slc32a1^tm2(cre)Lowl^ knock-in mice, which express Cre recombinase in inhibitory GABAergic neurons under control of the vesicular GABA transporter promoter, are referred to as VGAT^iCre 105^. All transgenic animals were backcrossed to C57BL/6 for several generations. Heterozygous Cre^+^ mice were used in experiments. Standard genotyping primers are available on the Jackson Lab website.

### Stereotactic injections and optic fiber implants

Conditional expression of target genes in Cre-containing neurons was achieved using recombinant adeno-associated viruses (AAVs) using the FLEX cassette or encoding a double-floxed inverted open reading frame (DIO) of target genes, as described previously^50^. For fiber photometry and *ex vivo* 2-photon calcium imaging experiments in the VTA, DAT^iCre^ mice were transduced with AAV1.CAG.FLEX.GCaMP6f.WPRE-SV40 (1.33 x 10^13^ GC/ml) from the UPenn viral core (Philadelphia, PA, a gift from the Genetically Encoded Neuronal Indicator and Effector Project (GENIE) and Douglas Kim; Addgene viral prep #100835-AAV1)^106^ or AAV8.CAG.FLEX.EGFP (3.1 x 10^12^ GC/ml, UNC vector core, Dr. Ed Boyden). For fiber photometry experiments in the mPFC, VGAT^iCre^ or Drd1^Cre^(FK150) mice were transduced with AAV1.CAG.FLEX.GCaMP6f.WPRE.SV40 (1.33 x 10^13^ GC/ml). For chemogenetic experiments in mPFC, Drd1^Cre^(FK150) mice were transduced with AAV2.hSyn.DIO.hM3Dq.mCherry (8.6 x 10^12^ GC/ml, Addgene viral prep #44361-AAV2, Dr. Bryan Roth)^107^ or AAV1.CBA.DIO.rM3Ds.mCherry.WPRE (4.3×10^13^ GC/ml, packaged by Vigene Biosciences). rM3Ds.mCherry sequence (Addgene #50458, Dr. Bryan Roth) was cloned into pAAV-CBA-DIO backbone (Addgene #81008, Dr. Bernardo Sabatini)^108^ between AscI and NheI restriction sites. For chemogenetic experiments in VTA, DAT^iCre^ mice were transduced with a custom built AAV1.CBA.DIO.hM4Di.mCherry (1.28 × 10^13^ GC/ml, Vigene Biosciences, Rockville, MD, plasmid a gift from Dr. Bernardo Sabatini)^108^. For glutamate uncaging-evoked spinogenesis experiments, AAV8.CAG.FLEX.EGFP (3.1 × 10^12^ GC/ml, UNC vector core, Dr. Ed Boyden) was co-injected with AAV1.hSyn.Cre.WPRE.hGH (1 × 10^10^, UPenn viral core, Dr. James M. Wilson, unpublished) to achieve sparse expression in mPFC pyramidal neurons of C57BL/6 or Drd1^ff^ mice. To achieve cell-autonomous inhibition of PKA via expression of PKIα in mPFC, AAV1.FLEX.PKIα.IRES.nls.mRuby2 (1.35×10^13^ GC/ml, packaged by Vigene Biosciences, Addgene #63059)^109^ was co-injected with AAV1.hSyn.Cre.WPRE.hGH and AAV8.CAG.FLEX.EGFP. For optogenetics experiments, DAT^iCre^ mice were transduced with AAV1.EF1a.DIO.hChR2(H134R).mCherry (1.2 × 10^13^ GC/ml) or AAV1.EF1a.DIO.hChR2(H134R).eYFP (3.55 × 10^13^ GC/ml), acquired from Dr. Karl Deisseroth (Addgene viral prep #20297/20298-AAV1), or AAV8.CAG.FLEX.EGFP for controls. For fiber photometry of dLight1.1 transients, custom built AAV1.CBA.DIO.dLight1.1.WPRE.hGh (1.22 × 10^13^ GC/ml, packaged by Vigene Biosciences) was used. dLight1.1 sequence was PCR amplified from pCMV.dLight1.1 (Addgene #111053, Dr. Lin Tian)^56^ using Q5 polymerase (primers: 5’-aaaagctagcatgaagacgatcatcgccctgagc-3’ and 5’-aaaaaggcgcgcctcaggttgggtgctgaccg-3’) and subcloned into pAAV.CBA.DIO (Addgene #81008)^108^ backbone between AscI and NheI restriction sites. For retrograde labeling, mice were intracranially injected with Red Retrobeads™ (Lumafluor Inc.).

Neonatal viral transduction was carried out to minimize invasiveness and increase surgical efficiency^49,50,110–112^. P3-6 mice were cryoanesthetized, received ketoprofen for analgesia, and were placed on a cooling pad. Virus was delivered at a rate of 100 nl/min for up to 150-200 nl using an UltraMicroPump (World Precision Instruments, Sarasota, FL). Medial prefrontal cortex (mPFC) was targeted in the neonates by directing the needle immediately posterior to the eyes, 0.3 mm from midline, and 1.8 mm ventral to skin surface. Ventral tegmental area (VTA) was targeted in the neonates by directing the needle approximately ±0.2 mm lateral from Lambda and 3.8 mm ventral to skin surface. Coordinates were slightly adjusted based on pup age and size. Following the procedure, pups were warmed on a heating pad and returned to home cages.

For intracranial AAV or retrobead injections, P25-30 mice were anesthetized with isoflurane (3% for induction value, 1.5-2% for maintenance) or ketamine:xylazine (100:12.5 mg/kg b.w.), received ketoprofen for analgesia, and were placed on a small animal stereotax frame (David Kopf Instruments, Tujunga, CA). AAVs or retrobeads were delivered unilaterally through a pulled glass pipette at a rate of 100 nl/min using an UltraMicroPump (World Precision Instruments, Sarasota, FL). Injection coordinates for VTA, 2.8 mm posterior to bregma, 0.4 mm lateral, and 4.3 - 4.5 mm below the pia; for mPFC, 2.0 mm anterior to bregma, 0.4 mm lateral, and 1.3 - 1.6 mm below the pia. Pipettes were held at the injection location for 15 min following AAV or retrobead release. Coordinates were slightly adjusted based on mouse age and size. Mice recovered for 7-9 days and > 14 days following retrobead and AAV injections, respectively.

For photometry fiber placement in the VTA or mPFC, mice were implanted with a 400 µm diameter 0.48 NA single mode optical fiber (Doric lenses, Quebec City, QC, Canada) directly above the VTA at -2.8 mm (AP); +0.4 mm (ML); 4.3 - 4.5 mm (DV), mPFC +2.0 mm (AP); +0.4 mm (ML); 1.3 - 1.6 mm (DV), four to six weeks after viral transduction. For optogenetics in the mPFC, custom-made optical fibers were used (2 mm stub length, 200 µm diameter, 0.5 NA, FP200ERT, Thorlabs). Coordinates for mPFC fiber placement were +2.0 mm (AP), +0.4 mm (ML), and 1.3 - 1.6 mm (DV). For bilateral cannulation in mPFC: +2.2 mm (AP); 1.2 mm (DV), with center to center distance of 0.8 mm. Dummy cannulae with projections of 0.2 mm over the guide cannulae (2.5 mm) were inserted after implantation. Internal cannulae with projections of 0.5 mm were used for liquid delivery. Behavioral experiments were conducted 7-12 days after implantation. Real-time photometry recording was performed during optical fiber implant above the VTA, for optimal targeting. When the fiber tip approached the VTA region, a continuous increase of fluorescence intensity was observed. The final position of implantation was determined by the cessation of further increases in fluorescence intensity.

### Behavior assays

#### Aversive learning

P40-60 mice were used for behavioral assays with fiber photometry, optogenetics and chemogenetics experiments. P25 - 40 mice were used for spinogenesis assays with behavioral manipulations. The strong learned helplessness procedure consisted of two induction sessions (1 session per day; 360 inescapable foot shocks per session; 0.3 mA, 3 sec; random 1-15 sec inter-shock intervals). Active/Passive Avoidance Shuttle Boxes from MazeEngineers (Boston, MA) were used for the experiment. To assess the degree of aversive learning, test sessions (30 escapable foot shocks per session; 0.3 mA, 10 sec; random 5-30 sec inter-shock intervals) were conducted before induction, 24 hrs after the last induction session, and following pharmacological or optogenetic manipulations. The testing was performed in a shuttle box (18 × 18 × 20 cm) equipped with a grid floor and a door separating the two compartments. No conditioned stimulus was delivered either before or after the shocks. Escapes were scored when the animal shuttled between compartments during the shock. Escape latency was measured as the time from the start of the shock to the escape. The shock automatically terminated when the animal shuttled to the other compartment. Failures were scored when the animal failed to escape before the shock end. All behavioral assays were conducted during the active phase of the circadian cycle. Schematics with mice were made using BioRender.com.

To evaluate the effect of ketamine on learned helplessness behavior, a single dose of ketamine (10 mg/kg b.w., i.p.) was given 48 hrs after the last induction session, and the test session performed 4 hrs later. For chemogenetic activation in mPFC, 48 hrs after the last induction session, Clozapine N-oxide (CNO) was administered i.p., followed by test sessions 2 hrs and 4 hrs later. For chemogenetic inhibition in VTA, the first CNO dose (3 mg/kg, i.p.) was co-administered with ketamine 48 hrs after the last induction, followed by test sessions 4 and 24 hrs later. Then, immediately following the last test session ketamine was administered alone, followed by test sessions 4, 24, and 72 hrs later. For optogenetic activation of DA terminals in the mPFC, 48 hrs after the last induction session a test session was performed with optogenetic stimulation. Bursts were not time-locked to behavior and consisted of ten 20 ms long pulses at 20 Hz, with 500 ms long burst duration and 10 sec inter-burst interval. Optical power of light at the tip of fiber was <9 mW/mm^2^. Light stimulation was applied in either compartment of the shuttle box.

For the weak aversive learning procedure, a single induction session (360 inescapable foot shocks; 0.3 mA, 3 sec; random 1-15 sec intershock intervals) was performed at the end of the 2^nd^ day. Six test sessions were performed (1 session per day; 100 escapable foot shocks per session; 0.3 mA, 3 sec; random 5-15 sec intershock intervals). A single dose of ketamine (10 mg/kg b.w., i.p.) was given 4 hrs before the 5^th^ test session. The final test session was performed 24 hrs after ketamine treatment. Age-matched controls were given six test sessions, in the absence of induction sessions or ketamine treatment.

#### Locomotion test

To assess the locomotor activity, mice were placed in the center of a plastic chamber (48 cm × 48 cm × 40 cm) or in the shuttle box in a dimly lit room. Mice explored the arena for 15 min, with video (30 fps) and photometry recording performed during the final 10 min.

### Local drug infusion

To inhibit hM4Di-expressing DA terminals by local CNO infusion in mPFC, we used internal cannulae (28 gauge, Plastics One, Roanoke, VA) with 0.5 mm long projection beyond the implanted guide-cannula, as described above. 1 mM CNO (1 µL per side) was administered over a 2 min long injection period using a 10 µl Hamilton syringe (Hamilton Company, Franklin, MA). Dummy cannulae with 0.2 mm projection were inserted back after the delivery of CNO. The first infusion was performed immediately before ketamine injection, and the two subsequent infusions were performed 1.5 and 3 hrs after ketamine injection. Learned helplessness behaviors were assessed 4 and 24 hrs after ketamine treatment. For acute ketamine or ACSF local infusion in mPFC, ketamine (12.5 µg in 500 nl ACSF) or ACSF (500 nl) were delivered unilaterally through a pulled glass pipette at a rate of 100 nl/min using an UltraMicroPump (World Precision Instruments, Sarasota, FL). Injection coordinates for mPFC, 2.0 mm anterior to bregma, 0.4 mm lateral, and 1.3 - 1.6 mm below the pia.

### Fiber photometry

A modified version of Scanimage ^113^ was used for data acquisition. Signals were sampled at 1 kHz and downsampled to 250 Hz for time-locked and 10 Hz for non-time-locked analysis. Fluorescence signal was baseline adjusted in non-overlapping 100 sec windows as (*signal* − *median*(*signal*)) / *median*(*signal*), denoted as dF/F. Recordings were made during a subset of the sessions of learned helplessness and open field locomotion, as noted in the text.

### Acute slice preparation and electrophysiology

Coronal brain slice preparation was modified from previously published procedures^49,50,114^. Animals were deeply anesthetized by inhalation of isoflurane, followed by a transcardial perfusion with ice-cold, oxygenated artificial cerebrospinal fluid (ACSF) containing (in mM) 127 NaCl, 2.5 KCl, 25 NaHCO_3_, 1.25 NaH_2_PO_4_, 2.0 CaCl_2_, 1.0 MgCl_2_, and 25 glucose (osmolarity 310 mOsm/L). After perfusion, the brain was rapidly removed, and immersed in ice-cold ACSF equilibrated with 95%O_2_/5%CO_2_. Tissue was blocked and transferred to a slicing chamber containing ice-cold ACSF, supported by a small block of 4% agar (Sigma-Aldrich). Bilateral 300 µm-thick slices were cut on a Leica VT1000s (Leica Biosystems, Buffalo Grove, IL) in a rostro-caudal direction and transferred into a holding chamber with ACSF, equilibrated with 95%O_2_/5%CO_2_. Slices were incubated at 34°C for 30 min prior to electrophysiological recording and 2-photon calcium imaging. Slices were transferred to a recording chamber perfused with oxygenated ACSF at a flow rate of 2–4 ml/min at room temperature.

To assess the number of action potentials evoked by current injections, current clamp whole-cell recordings were obtained from neurons visualized under infrared DODT or DIC contrast video microscopy using patch pipettes of ∼2–5 MΩ resistance. Neurons received 500 ms long, -20 pA to 160 pA current injections (in 20 pA steps), randomized across 15 sec-long sweeps. To assess spontaneous firing rate of dopaminergic neurons in the VTA, cell-attached recordings were performed. Cell-attached recording electrode pipettes were filled with the internal solution for voltage clamp recordings to monitor spontaneous break in, with pipette resistance varying between 3 and 7 MΩ. Dopaminergic neurons were identified by the expression of mCherry in DAT^iCre^ mice. To activate ChR2-expressing fibers of VTA DA neurons in the mPFC, 2 ms-long light pulses (470 nm, ∼2 mW) at intervals of 30 sec were delivered at the recording site using whole-field illumination through a 60X water-immersion objective (Olympus, Tokyo, Japan) with a PE4000 CoolLED illumination system (CoolLED Ltd., Andover, UK). Voltage clamp whole-cell recordings were performed to measure light-evoked excitatory postsynaptic currents (EPSCs) in the presence of 10 µM gabazine. Recording electrodes contained the following (in mM): Current-clamp recordings: 135 K-gluconate, 4 KCl, 10 HEPES, 10 Na-phosphocreatine, 4 MgATP, 0.4 Na_2_GTP, and 1 EGTA (pH 7.2, 295 mOsm/L). Voltage clamp and cell-attached recordings: 120 CsMeSO_4_, 15 CsCl, 10 HEPES, 10 Na-phosphocreatine, 2 MgATP, 0.3 NaGTP, 10 QX314, and 1 EGTA (pH 7.2-7.3, ∼295 mOsm/L). Recordings were made using 700B amplifiers (Axon Instruments, Union City, CA); data were sampled at 10 kHz and filtered at 4 kHz with a MATLAB-based acquisition script (MathWorks, Natick, MA). Series resistance and input resistance were monitored using a 250 ms long, 5 mV hyperpolarizing pulse at every sweep, and experiments were started after series resistance had stabilized (<20 MΩ, uncompensated). Offline analysis of electrophysiology data was performed using Igor Pro (Wavemetrics, Portland, OR) and MATLAB (Mathworks, Natick, MA).

### Two-photon imaging and two-photon glutamate uncaging

Dendritic imaging and uncaging of MNI-glutamate for spinogenesis induction were accomplished on a custom-built microscope combining two-photon laser-scanning microscopy (2PLSM) and two-photon laser photoactivation, as previously described^49,50,64^. Two mode-locked Ti:Sapphire lasers (Mai Tai eHP and Mai Tai eHP DeepSee, Spectra-Physics, Santa Clara, CA) were tuned to 910 and 725 nm for exciting EGFP and uncaging MNI-glutamate, respectively. The intensity of each laser was independently controlled by Pockels cells (Conoptics, Danbury, CT). A modified version of Scanimage software was used for data acquisition^113^. For glutamate uncaging, 2.5 mM MNI-caged-L-glutamate (Tocris) was perfused into the slice chamber, and 725 nm light guided through a galvo scanhead was used to focally release the caging group. Secondary and tertiary dendritic branches were selected for dendritic imaging and spinogenesis induction. MNI-glutamate was uncaged near the dendrite (∼0.5 µm) at 2 Hz using up to forty 2 ms-long pulses. Images were continually acquired during the induction protocol at 1 Hz, and uncaging was stopped if a spinehead was visible before 40 uncaging pulses were delivered. Analysis was carried out on raw image stacks and z projections. For display purposes only, a subset of the 2-photon micrographs was processed using Candle^115^. A successful induction of new dendritic spine was scored when a protrusion from the dendrite in the uncaging location was observed. A newly generated dendritic spine had to satisfy the following criteria: *de novo* protrusion from the dendrite within 1 µm of the uncaging site; mean spine head fluorescence matching average fluorescence of spine heads on the parent dendrite; mean spine head fluorescence exceeding 20% of intensity in the parent dendrite. Changes in fluorescence intensity were profiled using line-scan analyses. For each animal, the probability of spinogenesis is represented as the fraction of successful induction trials out of all conducted trials within individual, (*P*(*spinogenesis*) = *number of succesful trials/total number of trials*).

Calcium imaging of GCaMP6f expressing DA neurons in acute brain slices of the VTA was done at 910 nm and sampled at 12 Hz. Spontaneous activity was imaged for 5 min in the baseline at 34°C, followed by a 5 min recording after *ex vivo* ketamine application (50 μM, with 15 - 40 min delay).

### Pharmacology

Pharmacological agents were acquired from Tocris (Bristol, UK) or Sigma-Aldrich (St. Louis, MO). *In vivo* injections included intraperitoneal and subcutaneous injections of ketamine (10 mg/kg, Vedco, St. Joseph, MO), SKF 83566 (10 mg/kg, Tocris), Clozapine N-oxide (3 mg/kg *in vivo*, 1 μM *in vitro*, Sigma-Aldrich). Ketamine (50 μM, Vedco), SR 95531 hydrobromide (gabazine, 10 μM, Tocris), SKF 81297 hydrobromide (SKF 81297, 1 μM, Tocris), and H-89 dihydrochloride (H-89, 10 μM, Tocris) were used in acute slice experiments, as noted.

### Tissue processing and immunohistochemistry

Mice were deeply anaesthetized with isoflurane and transcardially perfused with 4% paraformaldehyde (PFA) in 0.1 M phosphate buffered saline (PBS). Brains were post-fixed for 1-5 days and washed in PBS, prior to sectioning at 50-100 µm on a vibratome (Leica Biosystems). Sections were pretreated in 0.2% Triton X-100 for an hour at RT, then blocked in 10% bovine serum albumin (BSA, Sigma-Aldrich, ST Louis, MO):PBS with 0.05% Triton X-100 for two hours at RT, and incubated for 24-48 hrs at 4°C with primary antibody solution in PBS with 0.2% Triton X-100. On the following day, tissue was rinsed in PBS, reacted with secondary antibody for 2 hrs at RT, rinsed again, then mounted onto Superfrost Plus slides (ThermoFisher Scientific, Waltham, MA). Sections were dried and coverslipped under ProLong Gold antifade reagent with DAPI (Molecular Probes, Life Technologies, Carlsbad, CA) or under glycerol:TBS (9:1) with Hoechst 33342 (2.5µg/ml, ThermoFisher Scientific). Primary antibodies used in the study were rabbit anti-tyrosine hydroxylase (1:1000; AB152, Abcam, Cambridge, UK), chicken anti-GFP (1:2000; AB13970, Abcam, Cambridge, UK), rabbit anti-RFP (1:500, 600-401-379, Rockland, Limerick, PA), pCREB S133 (1: 5000, Abcam, ab32096), and rabbit anti-c-Fos (1:5000; Synaptic Systems, Goettingen, Germany). Alexa Fluor 488- or Fluor 647-conjugated secondary antibodies against rabbit or chicken (Life Technologies, Carlsbad, CA) were diluted 1:500. Whole sections were imaged with an Olympus VS120 slide scanning microscope (Olympus Scientific Solutions Americas, Waltham, MA). Confocal images were acquired with a Leica SP5 confocal microscope (Leica Microsystems). Depth-matched z-stacks of 2 µm-thick optical sections were analyzed in ImageJ (FIJI)^116,117^. For pCREB and c-fos quantification, every four adjacent z stack slices were combined, for a total of 6 µm thickness. mCherry signal was used to localize cell bodies of hM3Dq-expressing neurons. Laser intensity and all imaging parameters were held constant across samples, and the same threshold was applied for subtracting background immunofluorescence. c-fos^+^ or pCREB^+^ neurons were identified by an experimenter blind to the conditions.

### Quantitative fluorescence *in situ* hybridization

Quantitative fluorescence *in situ* hybridization (FISH) was conducted following previously published procedures^72,114^. Mice were deeply anesthetized by inhalation of isoflurane and decapitated. Brains were quickly removed and frozen in tissue-freezing medium on a mixture of dry ice and ethanol for 5 - 15 min prior to storage at 80°C. Brains were subsequently cut on a cryostat (Leica CM1850, Leica Biosystems) into 20 µm-thick sections, adhered to Superfrost Plus slides, and frozen at 80°C. Samples were fixed with 4% PFA in 0.1 M PBS at 4°C for 15 min, processed according to the manufacturer’s instructions in the RNAscope Fluorescent Multiplex Assay manual for fresh frozen tissue (Advanced Cell Diagnostics, Newark, CA), and coverslipped with ProLong Gold antifade reagent with DAPI (Molecular Probes). Enhanced green fluorescent protein channel 1 (*Egfp*) and dopamine Drd1a receptor channel 2 (*Drd1a*) probes were added to slides in combination, and Amp4-b fluorescent amplification reagent was used for all experiments. Sections were subsequently imaged on a Leica SP5 confocal microscope in four channels with a 40x objective lens at a zoom of 1.4 and resolution of 512 x 512 pixels with 1.0 µm between adjacent z sections. Images were taken across the entire population of mPFC *Egfp-* positive neurons in each brain section.

FISH images were analyzed using FIJI^116^. Briefly, every four adjacent z stack slices were combined, for a total of 3 µm thickness, in order to minimize missed colocalization, while decreasing false positive colocalization driven by signal from cells at a different depth in a similar x-y position. All channels were thresholded. Cellular ROIs were defined using the *Egfp*^+^ or retrobead^+^ channel information to localize cell bodies. FISH molecule puncta were counted within established cell boundaries. Whether a cell was considered positive for a given marker was determined by setting a transcript-dependent threshold of the number of puncta (e.g., over 5 puncta/soma for *Drd1a*^+^). These stringent parameters for co-localization and the challenges of quantifying low abundance receptor transcripts likely lead to underestimation of receptor-positive population sizes.

### Quantification of behavior

For evaluating locomotor behavior, Toxtrac^118^ was used to track the animal’s position, defined by its body center position, and quantify the distance travelled in each session. To detect motion onset/offset, movement was defined by the body center moving > 30 mm/sec for at least 0.5 sec. The associated Ca^2+^ transients were then aligned to transitions between motion start and stop times and averaged across all animals. In a subset of animals, contextual freezing behavior was evaluated in the learning or novel contexts with a single 5 min long exposure^119^ before the test session for aversive learning. Video was recorded and the mouse was tracked with Toxtrac. A freezing bout was defined as traveling <5 mm within a 3 sec bin; and the total time of freezing was expressed as % of the 5 min interval.

Three state transition models were constructed for a given condition and behavioral response, by counting the occurrence of each of the 3 possible 2-response sequences in all animals and then dividing by the total number of the given response. For example, an escape can be followed by another escape, a failure, or a premature trial. The probability of a failure following an escape was calculated by counting the number of failures that followed escapes and dividing it by the total number of escape trials. The graphs for the transition models were constructed using GraphViz^120^. The similarity between transition models was calculated as one minus the Frobenius norm (the Euclidean norm of a matrix) of the difference between models (1 − *norm*(*model*_*a*_ − *model*_*b*_)). To depict individual learning trajectories, starting at zero, trajectory value incremented by 1 for each escape trial, decremented by 1 for each failure trial, and kept constant for spontaneous transitions in the shuttle box.

### Quantification of fiber photometry data

The heatmap of Ca^2+^ transients was constructed by plotting single trial transients with signal amplitude depicted by color. The latency to peak was defined as the time from shock onset to the maximum Ca^2+^ transient value within 8 sec of shock onset. The negative and positive area under the curve (AUC) was calculated based on the direction of the peaks. Peaks were omitted from analyses if they were less than 5% of the distance from minimum to maximum dF/F or less than 3 times the standard deviation of the baseline period. The baseline was chosen by using the mean dF/F of the 5 sec segment before the shock onset. For behavioral response-specific Ca^2+^ transients, the transients for each behavioral response were averaged within a single subject, and then these data were averaged across all animals. The distance between escape and failure traces for each animal was computed as the Euclidean norm of the difference between the average of all their Ca^2+^ transients on escape trials and the average of all their Ca^2+^ transients on failure trials (||*mean*(*escape trials*) − *mean*(*failure trials*)||_2_).

### Quantification of dendritic spine density

Sections of mPFC were either examined with a custom-built 2PLSM or a Leica SP5 confocal microscope (Leica Microsystems). Distal apical dendritic segments were selected for analysis. For each dendritic segment, dendritic spines protruding on both sides of the dendrite were marked using a 3D reconstruction system Neurolucida 360 (MBF Bioscience, Williston, VT). Six to eight z stacks (0.3 µm between each stack), at 0.07 µm lateral pixel size, were used for reconstruction. Dendritic spine density was averaged from eight to twelve dendritic segments for each animal.

### Quantification of two-photon Ca^2+^ imaging

ROIs of dopaminergic neuron somata were defined manually. Raw fluorescence intensity for all frames during 5 min recording sessions was extracted using FIJI^116^. For each neuron, the power spectral density of its baseline adjusted fluorescence was computed using Welch’s method^121^ with a Hann window and 50% overlap.

### Statistical analyses

Group statistical analyses were done using GraphPad Prism 7 software (GraphPad, LaJolla, CA). For N sizes, the number of trials or cells recorded, as well as the number of animals are provided. All data are expressed as mean ± SEM, or individual plots. Probabilities are expressed as aggregate probabilities within individuals. For two-group comparisons, statistical significance was determined by two-tailed Student’s t-tests. For multiple group comparisons, one-way analysis of variance (ANOVA) tests were used for normally distributed data, followed by post hoc analyses. For non-normally distributed data, non-parametric tests for the appropriate group numbers were used. Pearson regression was used to detect the correlation between two groups of data. P < 0.05 was considered statistically significant.

## Extended Data Figures and Figure Legends

**Extended Data Figure 1.**
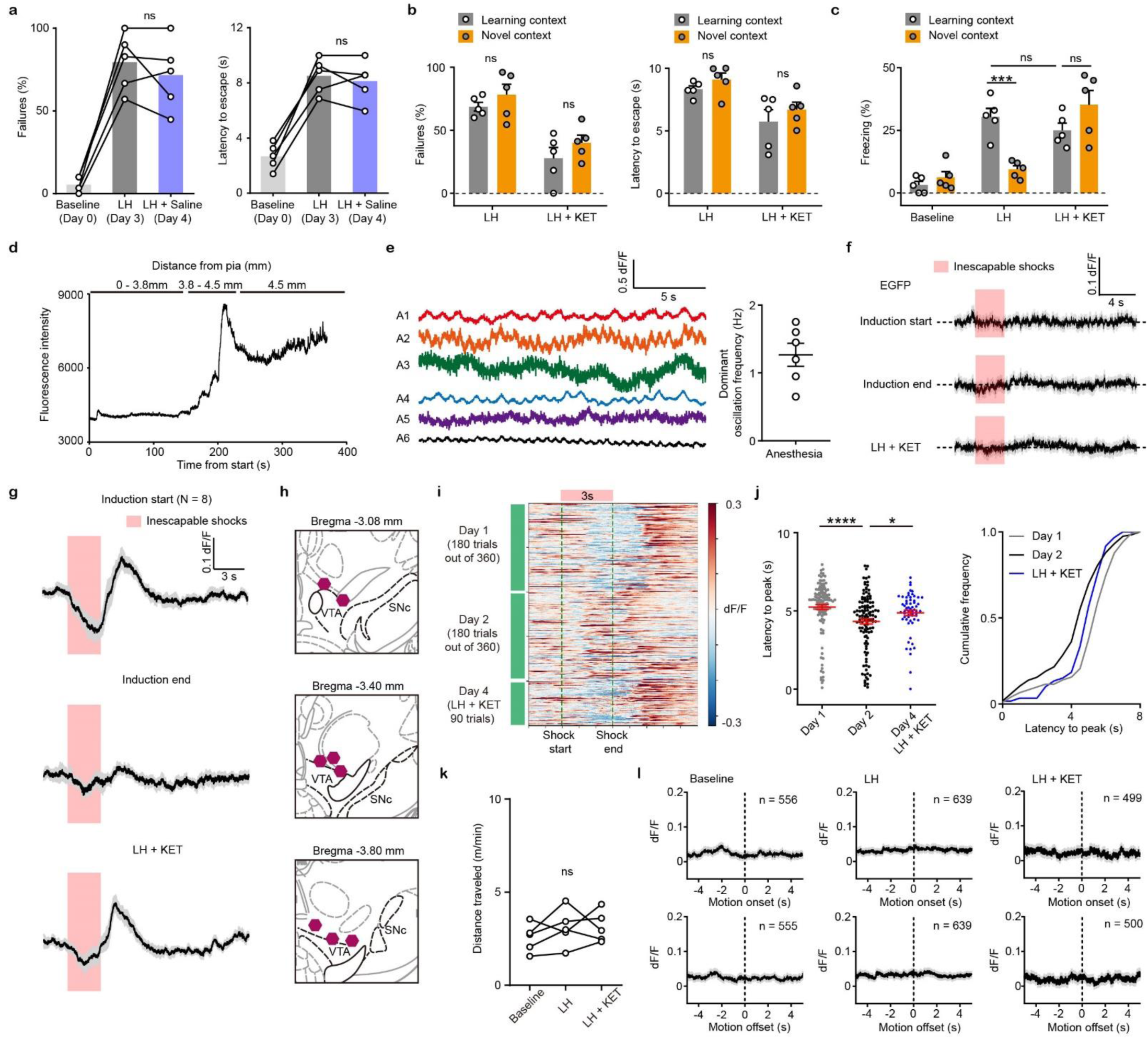
VTA DA neuron responses to anesthesia, foot shock, and motion transitions. (a). Left, summary data showing the percentage of failures to escape an escapable aversive shock across phases of learning (Baseline, LH, and LH + Saline). Right, same as left, but for latency to escape. n = 5 animals. One-way ANOVA, Sidak’s multiple comparison test, LH vs LH + KET, % Failures, p = 0.5299, latency to escape, p = 0.7027. (b). Left, summary data showing the percentage of failures to escape an escapable aversive shock in learning and novel contexts (LH, and LH + KET). Right, same as left, but for latency to escape. n = 5 animals. Two-way ANOVA, Sidak’s multiple comparison test, learning vs novel contexts, % Failures, LH, p = 0.5669, LH + KET, p = 0.3924. Latency to escape, LH, p = 0.6409, LH + KET, p = 0.5089. (c). Summary data showing the percentage of freezing behavior in learning or novel contexts across phases of learning (Baseline, LH, and LH + Saline). n = 5 animals/group. Two-way ANOVA, Sidak’s multiple comparison test, learning vs novel contexts, Baseline, p = 0.8892; LH, p = 0.0004; LH + KET, p = 0.1041. (d). Photometry-guided fiber implantation into the VTA. Recording began when the fiber traversed the pia. A gradual increase of fluorescence intensity was observed as the fiber tip approached GCaMP6f-expressing neurons, 3.8 – 4.5 mm from the pia, followed by a sharp signal increase and stabilization close to the VTA. (e). Oscillation of VTA DA Ca^2+^ transients under anesthesia. Approximately 1 - 2 Hz oscillations were observed in every animal. Summary data show the dominant oscillation frequencies. (f). Average neural activity-independent fluorescence transients illustrated for one EGFP-expressing animal in response to inescapable foot-shocks across learning phases. Traces are aligned to shock start time (20 trials/animal, n = 3 animals). (g). Average Ca^2+^ transients (mean ± SEM) in response to foot shocks for all animals across learning phases. Traces are aligned to shock start time (20 trials/animal, 8 animals). (h). Atlas locations showing fiber placements for data in Figures 1e and Extended Data Figure 1g. (i). Heatmap of single trial Ca^2+^ traces across learning in one mouse. Pink rectangle and green dashed lines mark the timing of shock stimuli. Colormap of dF/F, blue -0.3, red 0.3, saturated for values outside this range for illustration purposes. (j). Left, latencies from shock onset to dF/F peak (averages of sequential bins of 10 traces) for subjects across learning days. Right, cumulative frequency distribution of latencies to peak. n = 8 animals, one-way ANOVA, F (2, 258) = 9.387, p = 0.0001. Sidak’s multiple comparison test, Day 1 vs Day 2, p = 0.0001, Day 2 vs LH+KET, p = 0.0342. (k). Open field locomotion (m/min) for n = 5 mice across learning phases. Repeated measures one-way ANOVA, F (1.038, 4.153) = 1.479, p = 0.2910. (l). Average Ca^2+^ transients around motion onset and offset for mice in the Baseline, LH, and LH + KET conditions. Dashed black line marks motion onset/offset. The number of onsets/offsets, as noted. n = 5 animals. *p < 0.05, *** p < 0.001, **** p < 0.0001. Error bars reflect SEM.

**Extended Data Figure 2.**
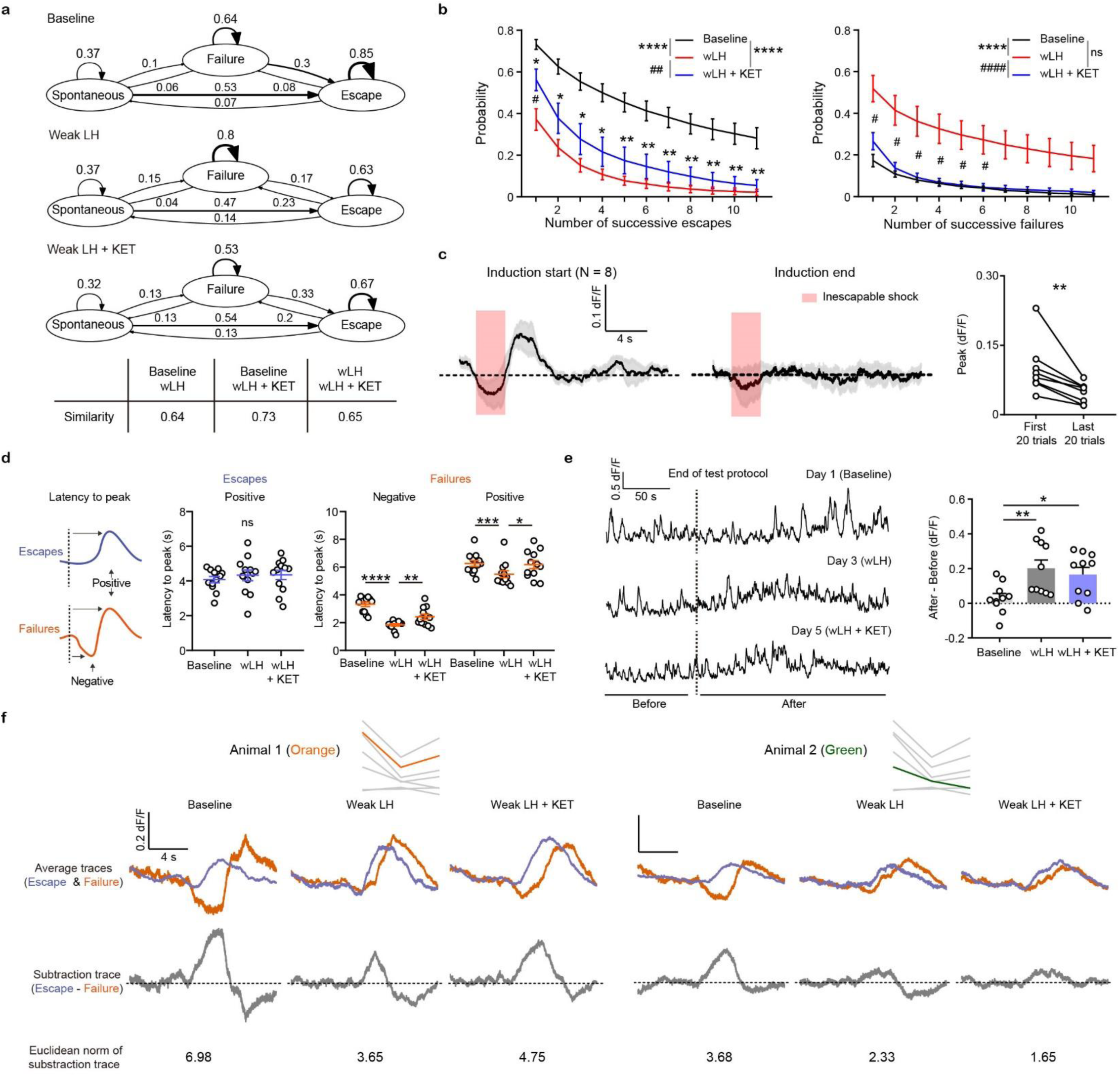
Characterization of behavioral sequences and VTA DA responses in weak learned helplessness (wLH) paradigm. (a). Graphs depicting transition probabilities between behavioral responses during baseline (top), wLH (middle), and wLH + KET (bottom). Arrow sizes are proportional to transition probability, noted numerically. Table, Frobenius norms of differences between graphs for each graph pair. n = 7 animals, 1400 trials/condition. (b). Summary data showing the probability of successive responses of different lengths (1-11) for escape (left) and failure (right) across learning states. Two-way ANOVA, Sidak’s multiple comparison test. Escape main effect, Baseline vs wLH and Baseline vs wLH + KET, p < 0.0001, wLH vs wLH + KET, p = 0.0031. Failure main effect, Baseline vs wLH and wLH vs wLH + KET, p < 0.0001, Baseline vs wLH + KET, p = 0.3291. For comparison within specific length of responses, * p < 0.05 ** p < 0.01 vs Baseline, # p < 0.05 vs wLH, n = 7 animals. (c). Left, average Ca^2+^ transients in response to foot shocks during wLH induction (mean ± SEM). Traces are aligned to shock start time (20 trials/animal, 8 animals). Right, quantification of peak Ca^2+^ transient amplitude of first and last 20 trials during induction. Two-tailed paired t-test, p = 0.009. (d). Left, schematic illustration for the measured variables. Middle, summary data for latency to peak of the positive Ca^2+^ transient peaks in escapes across learning phases. Right, same but for both positive and negative peaks in failures, as shown in the schematic. n = 12 animals, Escapes, positive peak, repeated measures one-way ANOVA, F (1.902, 20.92) = 0.622, p = 0.5388. Failures, negative peak, repeated measures one-way ANOVA, F (1.784, 19.62) = 36.23, p < 0.0001, Holm-Sidak’s multiple comparison test, Baseline vs wLH, p < 0.0001, wLH vs wLH + KET, p = 0.0025. Positive peak, repeated measures one-way ANOVA, F (1.301, 14.31) = 5.369, p = 0.0286, Holm-Sidak’s multiple comparison test, Baseline vs wLH, p = 0.0003, wLH vs wLH + KET, p = 0.0446. (e). Increase of overall fluorescent intensity after the end of wLH procedure across learning phases. Left, representative traces from the same subject in each condition. Right, summary data for post-protocol increases in fluorescent intensity. One-way ANOVA, F (2, 27) = 5.233, p = 0.012, Sidak’s multiple comparison test, Baseline vs wLH, p = 0.0097, Baseline vs wLH + KET, p = 0.044, n = 10 animals. (f). Example trace distance calculation for 2 animals. Top row, individual animals marked on the trace distance graph from Figure 2f. Second row, average traces of escapes and failures across learning states. Third row, difference between the average escape and failure traces. Bottom row, Euclidean norm of subtraction traces. *p < 0.05, ** p < 0.01, *** p < 0.001, **** p < 0.0001. ^#^p < 0.05, ^##^p < 0.01, ^####^p < 0.0001. Error bars reflect SEM.

**Extended Data Figure 3.**
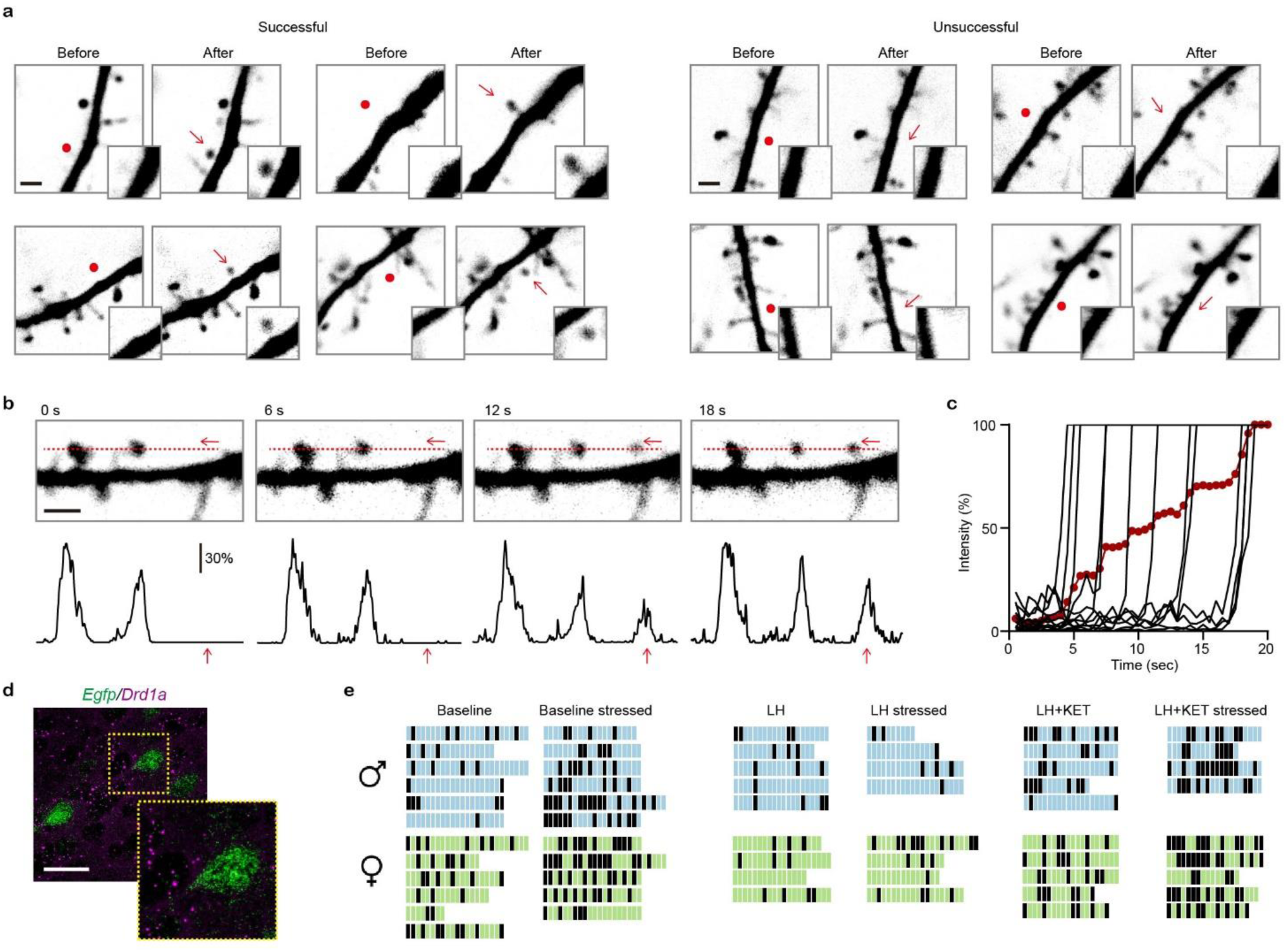
*De novo* glutamate-induced spinogenesis on pyramidal neurons in mPFC. (a). Example 2PLSM images of successful and unsuccessful induction trials of *de novo* spinogenesis. Red circles, uncaging sites. Inset, close up images of local dendritic segments before and after glutamate uncaging. Scale bar, 2 µm. (b). Top, time-lapse images of spine formation during glutamate uncaging (40 pulses, 2 Hz). Red arrow, uncaging spot and nascent spine. Line-scan analysis was performed, as noted by the red dashed line. Bottom, fluorescence intensity profiles from the line-scan across pre-existing spines and uncaging spot. Scale bar, 2 µm. (c). Time course of individual trials (2 Hz) and average (red dots) fluorescence intensity changes across a series of successful trials during glutamate uncaging. (d). Fluorescence *in situ* hybridization (FISH) image showing the absence of *Drd1a* mRNA expression (purple) in *Egfp* mRNA expressing mPFC cells (green). Inset, close up of a single neuron. Scale bar, 50 µm (e). Binary grid plots illustrating sequences of spinogenesis induction outcomes for different mice in all conditions in Figure 5l, separated by sex. Black, successful evoked spinogenesis trial; blue or green, unsuccessful trial for males or females, respectively.

**Extended Data Figure 4.**
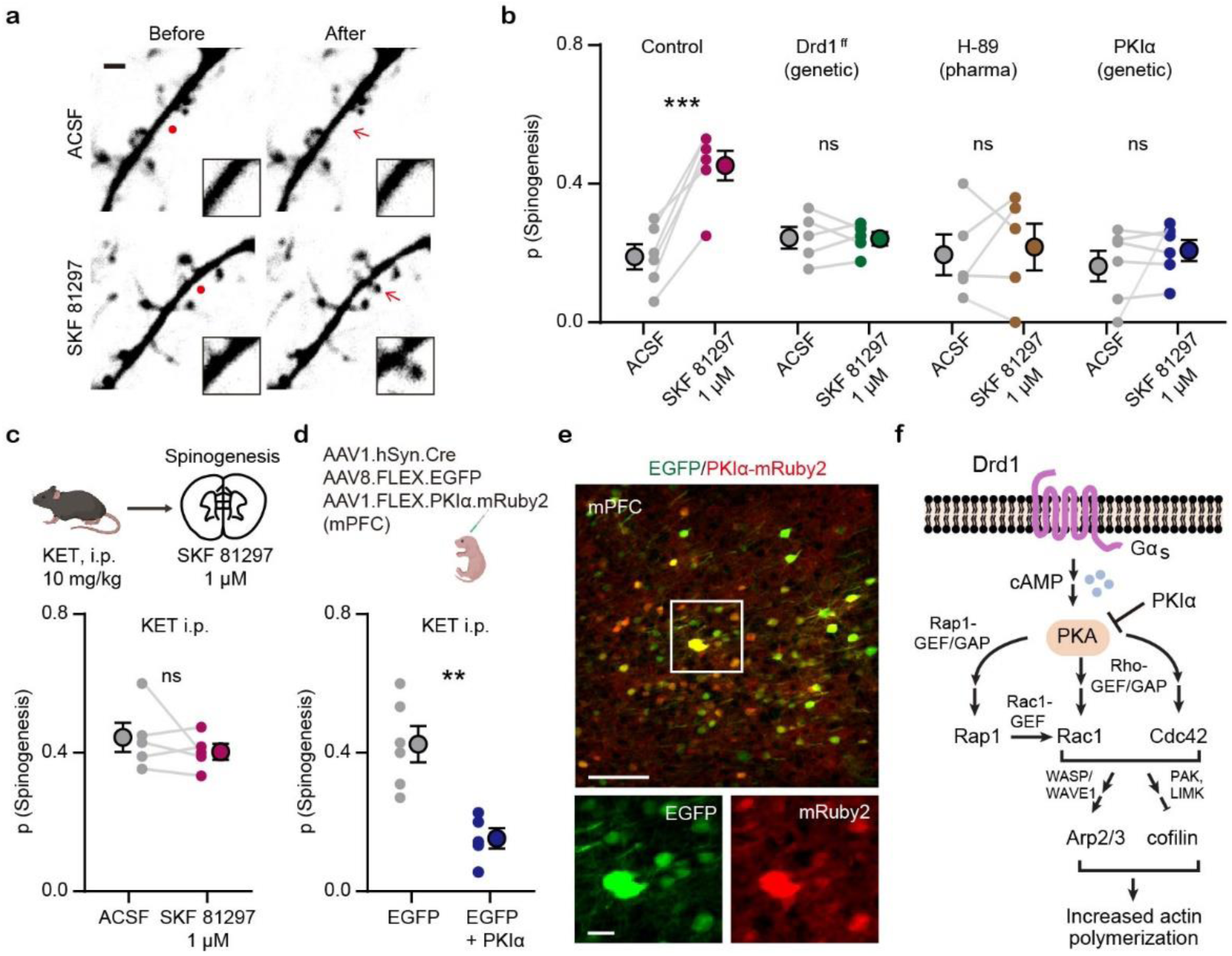
Drd1 activation promotes glutamate-induced spinogenesis in mPFC pyramidal neurons through PKA signaling. (a). Example 2PLSM images of *de novo* spinogenesis trials with ACSF or 1 µM SKF 81297. Red circles, uncaging sites. Black rectangle, close up images of local dendritic segments before and after glutamate uncaging. Scale bar, 2 µm. (b). Probability of glutamate-evoked spinogenesis on deep layer mPFC neurons in brain slices with or without bath application of 1 µM SKF 81297. Slices were treated with 10 µM H-89 or collected from mice with genetic manipulation of GFP expressing pyramidal neurons (Drd1 ff or PKIα). Each small circle, aggregate probability of evoked spinogenesis from a single experiment. Large circles, group data. Paired two-tailed t test, ACSF vs SKF 81297, Control, p = 0.0007; Drd1 ^ff^, p = 0.9249; H-89, p = 0.7351; PKIα, p = 0.4; n = 5 - 6 experiments/group. (c). Top, schematic illustrating glutamate-evoked spinogenesis assay in slices from mice pre-treated with ketamine (10 mg/kg, i.p.). Bottom, probability of glutamate-evoked spinogenesis on deep layer mPFC neurons in brain slices with or without bath application of 1 µM SKF 81297. Paired two-tailed t test, ACSF vs SKF 81297, p = 0.3745. (d). Top, schematic illustrating triple viral transduction strategy for PKIα expression. Bottom, probability of glutamate-evoked spinogenesis in deep layer mPFC neurons in mice with or without PKIα expression, injected with ketamine (10 mg/kg, i.p.). Unpaired two-tailed t test, GFP vs GFP + PKIα, p = 0.0020. (e). Top, colocalization of PKIα-mRuby2 in EGFP-expressing mPFC neurons. Bottom, close up images of EGFP and mRuby2 signals. Scale bar, 100 µm and 20 µm. (f). Schematic of simplified signaling pathways downstream of Drd1-PKA involved in actin remodeling in dendritic spines. **p < 0.01, *** p < 0.001. Error bars reflect SEM.

**Extended Data Figure 5.**
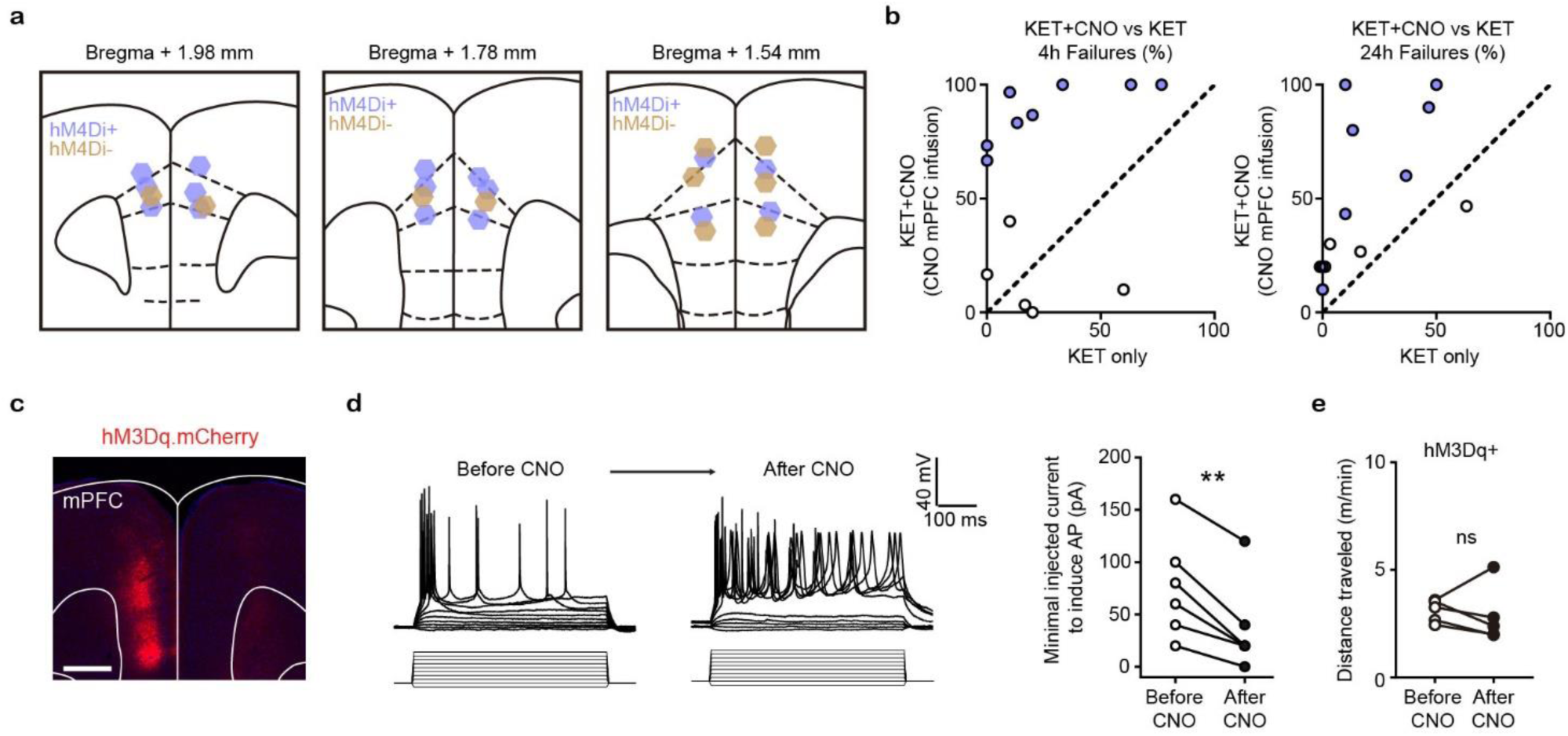
Local inhibition of DA terminals in mPFC blocks the behavioral effect of ketamine. (a). Atlas locations showing cannula placements for data in Figure 6c. (b). Within subject summary data for behavioral responses (Figure 6c) after ketamine treatment, compared with ketamine + CNO (mPFC), 4 hrs and 24 hrs after treatment, n = 5 - 8 animals. (c). Epifluorescent image of hM3Dq-mCherry expression in mPFC. Scale bar, 500 μm. (d). Whole-cell recording of action potentials evoked by different amplitude current injections in one mCherry^+^ mPFC neuron before, during, and after bath application of 1 µM CNO. Left, example traces. (a). Right, summary data for minimal injected current sufficient to evoke action potential firing before and after CNO application. n = 6 cells from 2 animals, two-tailed paired t-test, p = 0.0028. (e). Total distance traveled per minute in an open field locomotion assay for hM3Dq^+^ animals before and after CNO treatment. Two-tailed paired t-test, p = 0.6289, n = 5 animals.

**Extended Data Figure 6.**
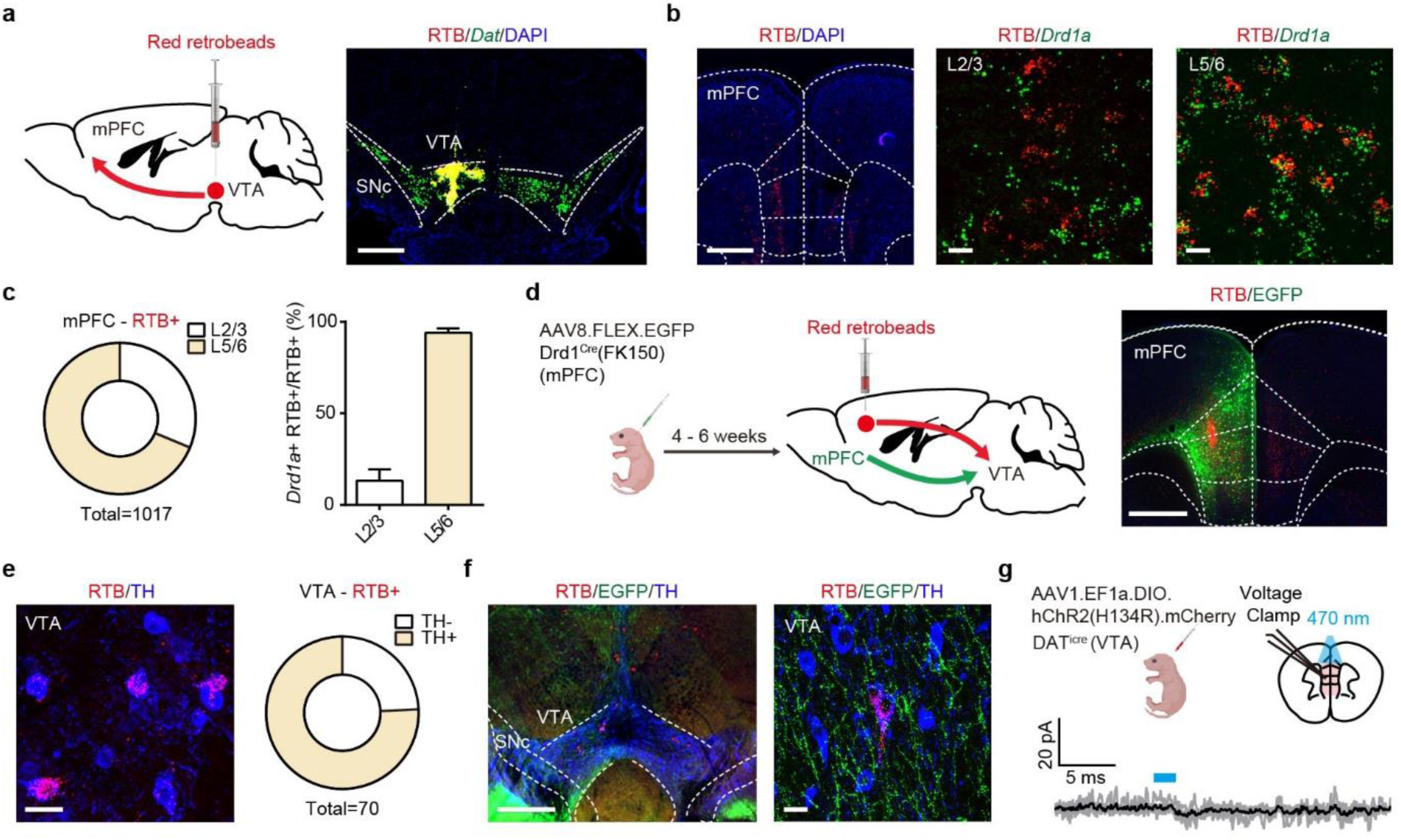
Recurrent loops link DA-sensing mPFC and VTA DA neurons. (a). Left, schematic illustrating retrograde labeling strategy. Right, coronal image of retrobead (RTB) injection site. Red, retrobeads; green, *Dat* mRNA; blue, DAPI. Scale bar, 500 μm. (b). Left, retrobead fluorescence in mPFC (atlas overlay, dashed line). Middle and right, colocalization of RTB and *Drd1* mRNA in superficial and deep layers of mPFC. Scale bars, 500 μm (left), 20 μm (middle and right). (c). Left, the proportional distribution of RTB^+^ neurons across superficial and deep cortical layers. Right, quantification of the percentage of *Drd1a*^*+*^ cells among RTB^*+*^ cells across layers. n = 2 animals, 31.3% RTB^+^ neurons in layer 2/3, 68.7% RTB^+^ in layer 5/6. Among those RTB^+^ neurons, 13.2 ± 6.2% are *Drd1*^*+*^ in layer 2/3, 94.1 ± 2.5 are *Drd1*^*+*^ in layer 5/6. (d). Left, schematic illustrating retrograde labeling with viral transduction strategy. Right, a coronal image showing RTB signal (red) and the expression of EGFP (green) in the mPFC. Scale bar, 500 μm. (e). Left, example image showing the colocalization of RTB (red) and TH (blue) in the VTA. Right, the proportional distribution of TH^+^ neurons among RTB^+^ cells in the VTA. Scale bar, 20 μm. n = 4 animals, 75.7% TH^+^. (f). Left, example image of EGFP^+^ terminals in the VTA and SNc (red, RTB; green, EGFP; blue, TH). Right, a close up image illustrating EGFP^+^ terminals surrounding TH^+^/RTB^+^ neurons. Scale bars, 500 μm, 20 μm. (g). Top, schematic illustrating viral transduction strategy and voltage-clamp recordings of mPFC pyramidal neurons. Bottom, summary data for voltage-clamp recordings of mPFC pyramidal neurons with optogenetic stimulation of VTA DA terminals (470 nm, 2 ms, ∼2 mW). Grey traces, optically evoked currents from individual neurons; black, average across cells. n = 5 cells from 2 animals.

**Extended Data Figure 7.**
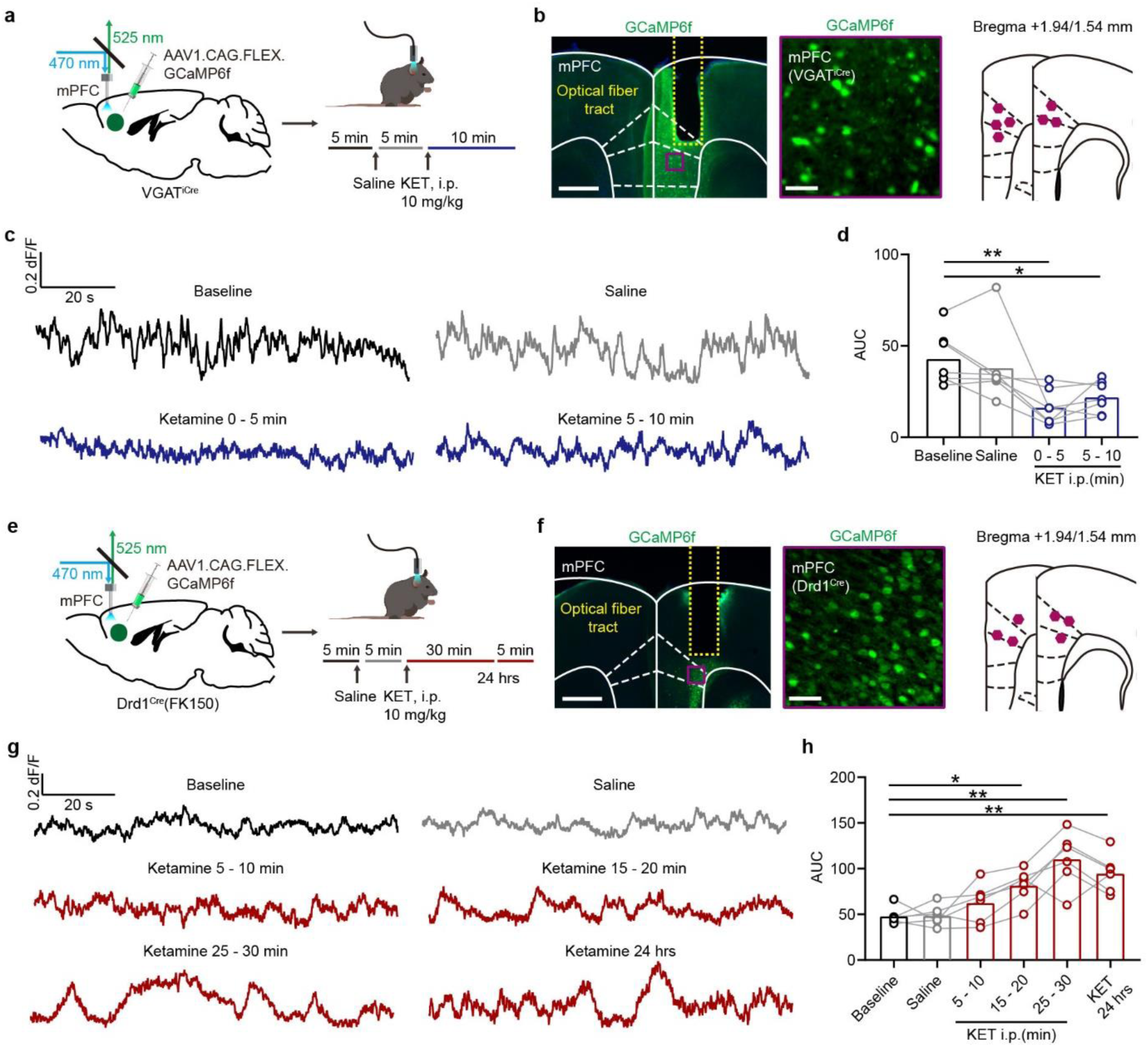
Activity of VGAT^+^ and Drd1^+^ neurons in mPFC after *in vivo* ketamine treatment. (a). Left, schematic for viral transduction of GCaMP6f in the mPFC with subsequent fiber implant in VGAT^iCre^ mice. Right, timeline of photometry recording with saline and ketamine treatment (10 mg/kg, i.p.). (b). Left, fiber placement illustration on a coronal section through mPFC, with a close up image of GABAergic neurons (white dashed lines, Paxinos atlas overlay; yellow dashed lines, fiber track). Green, GCaMP6f; blue, Hoechst nucleic stain. Scale bars: 500 μm and 50 μm. Right, atlas location of fiber placement for each subject. (c). Example traces showing Ca^2+^ transients of GABAergic neurons in mPFC from one mouse in the baseline condition, following a saline injection, and after ketamine (0 - 5 min and 5 - 10 min). Black, baseline; gray, saline; blue, ketamine. (d). Summary data showing area under the curve (AUC) of Ca^2+^ transients in 5 min bins across conditions (Baseline, Saline, 0 - 5, and 5 - 10 min after Ketamine). n = 7 animals, repeated measures one-way ANOVA, F (1.528, 9.167) = 8.836, p = 0.0098. Sidak’s multiple comparison test vs Baseline, Saline, p = 0.6653, KET 0 - 5 min, p = 0.0032, KET 0 - 5 min, p = 0.0325. (e). Left, schematic for viral transduction of GCaMP6f in the mPFC with subsequent fiber implant in Drd1^Cre^(FK150) mice. Right, timeline of photometry recording with saline and ketamine treatment (10 mg/kg, i.p.). (f). Left, fiber placement illustration on a coronal section through mPFC, with a close up image of Drd1^+^ (white dashed lines, Paxinos atlas overlay; yellow dashed lines, fiber track). Green, GCaMP6f; blue, Hoechst nucleic stain. Scale bars: 500 μm and 50 μm. Right, atlas location of fiber placement for each subject. (g). Example traces showing Ca^2+^ transients of Drd1^+^ neurons in the mPFC from one mouse in baseline, following saline treatment, and after ketamine treatment (5 - 10, 15 - 20, 25 - 30 min, and 24 hrs). Black, baseline; gray, saline; red, after ketamine. (h). Summary data showing area under the curve (AUC) of Ca^2+^ transients in 5 min bins across conditions (Baseline, Saline, 5 - 10, 15 - 20, 25 - 30 min, and 24 hrs after ketamine). n = 6 animals, repeated measures one-way ANOVA, F (2.498, 12.49) = 16.94, p = 0.0002. Sidak’s multiple comparison test vs Baseline, Saline, p > 0.9, KET 5 - 10 min, p = 0.3605, KET 15 - 20 min, p = 0.0131, KET 25 - 30, p = 0.0063, KET 24 hrs, p = 0.0031. *p < 0.05, ** p < 0.01.

**Extended Data Figure 8.**
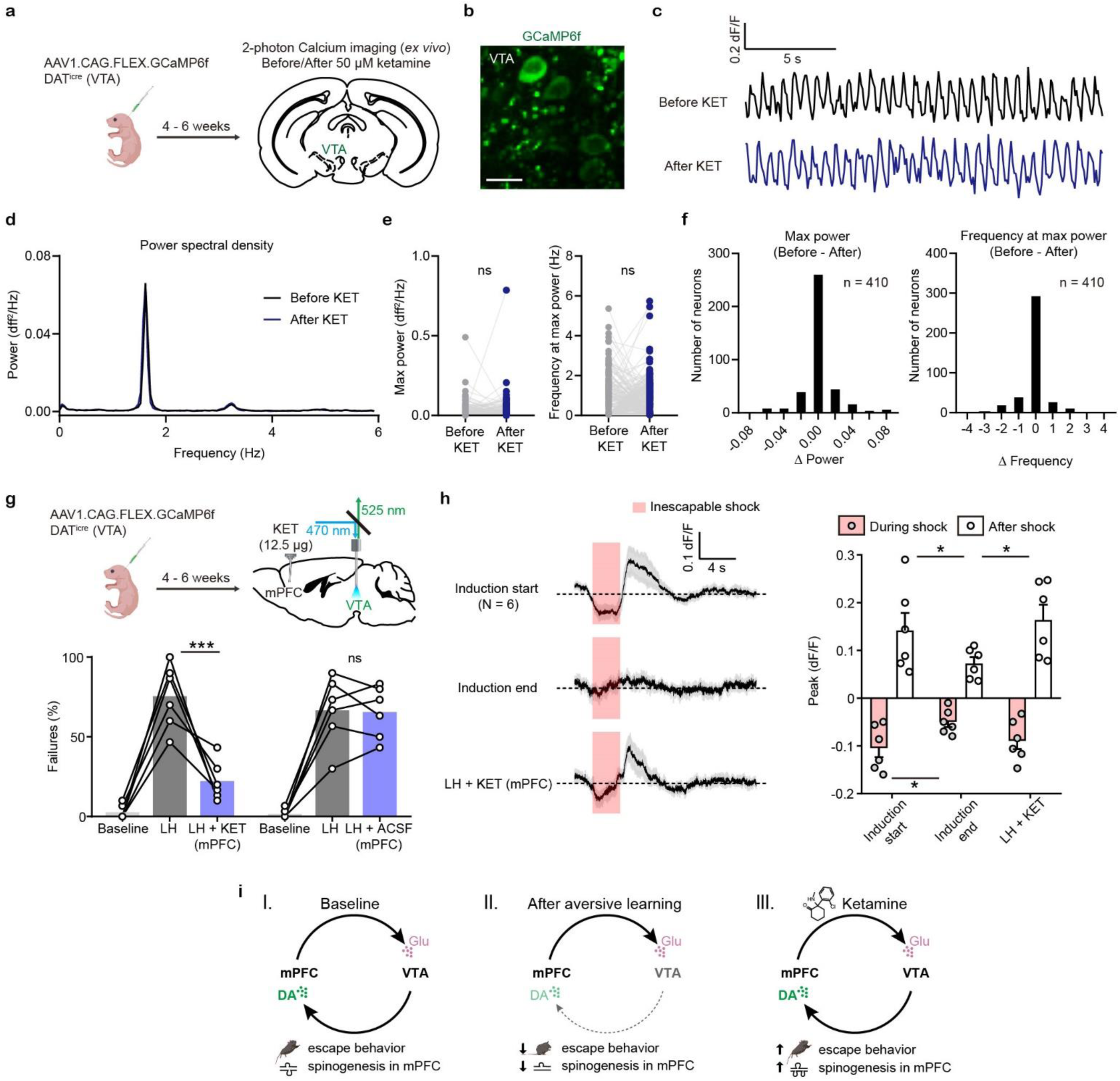
Local and circuit level effects of ketamine on VTA DA neurons. (a). Schematic illustrating viral transduction strategy and 2-photon Ca^2+^ imaging of VTA DA neurons in acute brain slices. (b). Example 2PLSM image of VTA DA neurons expressing GCaMP6f. Scale bar, 20 µm. (c). Spontaneous Ca^2+^ oscillations in one neuron before and after ketamine bath application (50 µM). Black, before ketamine; blue, after ketamine. (d). Power spectral density of Ca^2+^ transients for the neuron in (c). (e). Left, quantification of max power before and after ketamine treatment (n = 410 neurons). Right, quantification of frequency at max power. Paired two-tailed t test, before vs after ketamine, Max power, p = 0.8865, Frequency at max power, p = 0.3779. (f). Left, histogram showing the distribution of changes in max power after ketamine application. Right, same but for frequency at max power. n = 410 neurons. (g). Top, schematic for viral transduction and VTA photometry recording of Ca^2+^ transients with local ketamine delivery in mPFC. Bottom, summary data showing the percentage of failures after local infusion of ketamine or ACSF in mPFC. Two-way ANOVA, Sidak’s multiple comparison test, sLH vs sLH + KET, p < 0.0001, sLH vs sLH + ACSF, p = 0.9987, n = 6 animals. (h). Left, average Ca^2+^ transients (mean ± SEM) in response to foot shocks at the start of induction, at the end of induction, and following local ketamine infusion. Traces are aligned to shock start time (20 trials/animal, 6 animals). Right, quantification of peak Ca^2+^ transient amplitude during and after foot shock stimuli across conditions. Both positive and negative values are quantified. n = 6 animals, repeated measures one-way ANOVA, Holm-Sidak’s multiple comparison test, Peak: During shock, F (1.948, 9.739) = 5.547, p = 0.0252, Induction start vs Induction end p = 0.0433, Induction end vs LH+KET, p = 0.0823. After shock, F (1.468, 7.341) = 10.05, p = 0.0105, Induction start vs Induction end, p = 0.0462, Induction end vs LH+KET, p = 0.0147. (i). Schematic summary illustrating recurrent mPFC VTA circuits across learning states, with the observed effects on VTA DA activity, behavior, and mPFC plasticity. *p < 0.05, *** p < 0.001. Error bars reflect SEM.

